# A single dose of the catecholamine precursor Tyrosine reduces physiological arousal and decreases decision thresholds in reinforcement learning and temporal discounting

**DOI:** 10.1101/2022.02.09.479693

**Authors:** David Mathar, Mani Erfanian Abdoust, Deniz Tuszus, Tobias Marrenbach, Jan Peters

## Abstract

Supplementation with the catecholamine precursor L-Tyrosine might enhance cognitive performance, but overall findings are mixed. Here, we investigate the effect of a single dose of tyrosine (2g) vs. placebo on two key aspects of catecholamine-dependent decision-making: model-based reinforcement learning (2-step task) and temporal discounting, using a double-blind, placebo-controlled, within-subject design (n=28 healthy male participants). We leveraged drift diffusion models in a hierarchical Bayesian framework to jointly model participants’ choices and response times in both tasks. Furthermore, comprehensive autonomic monitoring (heart rate, heart rate variability, pupillometry, spontaneous eye-blink rate) was performed both pre- and post-supplementation, to explore potential physiological effects of supplementation. Across tasks, tyrosine consistently reduced participants’ RTs without deteriorating task-performance. Diffusion modeling linked this effect to attenuated decision-thresholds in both tasks and further revealed increased model-based control (2-step task) and (if anything) attenuated temporal discounting. On the physiological level, participants’ pupil dilation was predictive of the individual degree of temporal discounting. Tyrosine supplementation reduced physiological arousal as revealed by increases in pupil dilation variability and reductions in hear rate. Supplementation-related changes in physiological arousal predicted individual changes in temporal discounting. Our findings provide first evidence that tyrosine supplementation might impact psychophysiological parameters, and suggest that modeling approaches based on sequential sampling models can yield novel insights into latent cognitive processes modulated by amino-acid supplementation.

## Introduction

The market for dietary supplements that promise to boost cognitive functioning is consistently growing (Chopra et al., 2022). The freely-accessible catecholamine precursor L-Tyrosine has increasingly gained attention in (cognitive) neuroscience. Accumulating evidence suggests that tyrosine may boost cognitive functions that rely on catecholamine neurotransmission, specifically in demanding situations (Colzato et al., 2013; Deijen et al., 1999; Kühn et al., 2019; O’Brien et al., 2007). Tyrosine is converted to l-3,4-dihydroxyphenylalanine (L-DOPA) by the rate-limiting enzyme tyrosine hydroxylase and then catalyzed into dopamine. Downstream, β-hydroxylase can convert dopamine to noradrenaline (Molinoff & Axelrod, 1971). As tyrosine hydroxylase is saturated with tyrosine by only approximately 75%, dietary supplementation of tyrosine can potentially increase dopamine and noradrenaline synthesis and transmission (Moja et al., 1996).

Dopamine transmission strongly modulates reinforcement learning (RL) and value-based decision-making (Bayer & Glimcher, 2005; Bromberg-Martin et al., 2010; Schultz et al., 1997; Steinberg et al., 2013). Alterations in dopamine neurotransmission have been linked to maladaptive behaviors such as substance abuse and gambling disorder (Balodis et al., 2012; Kayser, 2019; Schultz, 2011). Learning from previous experience is essential for optimal behavior in volatile environments and is tightly linked to reward prediction errors signaled by dopaminergic midbrain neurons (Schultz et al., 1997). Reliance on stimulus-reward associations is referred to as model-free RL. In contrast, model-based RL is related to a computationally more demanding incorporation of a cognitive model of the environment to facilitate goal-directed action selection (Balleine & O’Doherty, 2010; Daw et al., 2011; Dolan & Dayan, 2013). The balance between model-free and model-based RL has been linked to striatal dopamine transmission (Deserno, 2015) and may constitute a trans-diagnostic marker across a range of psychopathologies (Culbreth et al., 2016; Gillan et al., 2016; Wyckmans et al., 2019). However, reported effects of a direct modulation of dopamine transmission on the balance of model-free and model-based RL remain mixed (Kroemer et al., 2019; Wunderlich et al., 2012).

Steep temporal discounting possibly delineates another trans-diagnostic marker with relevance for addiction-related disorders (Amlung et al., 2019; Lempert et al., 2019). temporal discounting refers to the trade-off between smaller but sooner (SS) and larger but later (LL) rewards (Peters & Büchel, 2011). Typically, both animals and humans prefer sooner over later rewards. The degree of temporal discounting is highly stable over time within individuals (Bruder et al., 2021; Enkavi et al., 2019; Kirby, 2009) and has been linked to dopaminergic function within the striatum (Joutsa et al., 2015). Parkinson’s disease patients show attenuated temporal discounting when tested on vs. off dopaminergic medication (Foerde et al., 2016). In line with this finding, de Wit et al. (2002) found that acute administration of D-amphetamine decreased temporal discounting in healthy volunteers. However, a later study did not replicate this effect (Acheson & de Wit, 2008). With respect to a direct modulatory role of dopamine, findings also appear mixed. Some studies have shown an increase in temporal discounting following moderate increases in dopamine transmission (Pine et al., 2010), and others showing the opposite (de Wit et al., 2002) or no modulatory effect (Petzold et al., 2019). Recently, Wagner et al. (2020) revealed attenuated temporal discounting following a single dose (2mg) of haloperidol, a dosage that might increase extra-synaptic striatal dopamine levels via blocking of presynaptic D2 auto-receptors.

Despite the relatively large body of work on pharmacological modulation of dopamine transmission and RL and temporal discounting, few studies have investigated possible modulatory effects of freely available tyrosine supplementation. So far, tyrosine supplementation has been shown to counteract decrements of cognitive performance in demanding situations in tasks related to dopamine transmission such as working memory (Hase et al., 2015; Jongkees et al., 2015). Several studies indicated that a single dose of tyrosine administration may improve a range of functions, including cognitive flexibility, inhibitory control, working memory, and reasoning (Colzato et al., 2013, 2014; Magill et al., 2003; Steenbergen et al., 2015). There is also evidence for effects of long-term tyrosine intake on cognitive performance, reflected in associations between daily tyrosine intake and working memory, episodic memory, and fluid intelligence (Kühn et al., 2019). However, no study to date reported effects of tyrosine administration on two potential trans-diagnostic markers, model-based RL and temporal discounting performance.

Here we examined the effect of a single dose of tyrosine (2g) on temporal discounting and model-based RL performance in a double-blind within-subject placebo controlled study. Importantly, we expanded upon previous work on supplementation effects in two ways. First, we utilized a combination of temporal discounting and reinforcement learning models with drift diffusion model based choice rules (Fontanesi et al., 2019; Pedersen et al., 2017; Peters & D’Esposito, 2020b; Shahar et al., 2019; Wagner et al., 2020). This approach yields a more comprehensive account of participants’ learning and decision-making behavior by decomposing observed RT distributions into latent underlying processes. Second, we comprehensively assessed effects of tyrosine supplementation on physiological proxy measures of catecholamine function, specifically spontaneous eye blink rate, pupil dilation, heart rate and variability of the latter two. Physiological measures were obtained both pre and post tyrosine/placebo administration. Results from these exploratory analyses revealed initial evidence for physiological effects of tyrosine supplementation and may inform future supplementation studies.

## Materials and methods

### Participants

Thirty male healthy volunteers (right-handed, non-smoking, with no history of psychiatric or neurological illness, no medication or drug use), participated in the study. We invited only male participants due to the potential impact of hormone levels on central tyrosine levels (Di Paolo, 1994; Diekhof & Ratnayake, 2016). All volunteers provided written informed consent and received a reimbursement of 10 € per hour for their participation. Performance dependent additional reimbursement is outlined in the task specific sections further below. The study was approved by the ethics committee of the German Psychological Association (DGPS). Two participants were excluded prior to data analysis, due to clinically significant depressive symptoms (BDI-II > 28), and a diagnosed psychiatric condition in the past.

### General procedure

A double-blind, placebo-controlled, randomized within-subjects design was employed. The study consisted of two testing sessions performed on two separate days with a 7 – 28 day interval in between. Participants were asked to refrain from drinking alcohol and eating protein-rich food on the evening before each testing day, to reduce possible interactions between tyrosine and other amino acids that might compete for amino acid transporter at the blood-brain barrier (Le Masurier et al., 2006). Participants were instructed to fast at least three hours before arriving and refrain from drinking anything except water. Testing started between 9 a.m. and 1 p.m. and lasted approximately 2.5 h.

Each testing day started with a five minute physiological baseline assessment (t0) including spontaneous eye blink rate, heart rate, and pupil dilation. After the baseline physiological recording session, tyrosine/placebo was provided to the volunteers (see Tyrosine / Placebo administration section below). This was followed by a 60 minutes break in which participants were free to rest and to read in a provided newspaper. Subsequently, participants underwent a second five minute physiological monitoring (t1) for exploratory analysis of possible tyrosine-related effects on physiological arousal. Then (75 minutes following tyrosine/PLC intake) participants performed the sequential RL task, followed by the temporal discounting task with a short resting break in between. The order of the tasks was fixed in both sessions. The sequential RL task is more challenging and time-consuming, and previous studies found an effect of tyrosine supplementation mainly in or after cognitively demanding situations (e.g. Colzato et al., 2013). In total, both tasks lasted for 45 minutes. The first testing day ended with a brief demographic and psychological screening (see below). At the end of the second testing day, participants received the payment and were asked to guess in which session they received tyrosine.

### Physiological data acquisition

For quantification of possible tyrosine-related effects on physiological arousal, we assessed three different measures of autonomic nervous system activity, spontaneous eye blink rate, heart rate, and pupil dilation for five minutes. Spontaneous eye blink rate is discussed as a proxy measure for central dopamine levels (Elsworth et al., 1991; Groman et al., 2014; Kaminer et al., 2011). Heart rate, pupil dilation and variability at rest in these measures are tightly linked to the interplay of sympathetic and parasympathetic afferents (Berntson et al., 1997; Bradley et al., 2008; Schneider et al., 2016) and noradrenaline transmission (Gordan et al., 2015; Murphy et al., 2014; Phillips et al., 2000; Preuschoff et al., 2011). In each of the two physiological assessments per testing day participants were seated in a shielded, dimly lit room 0.6m from a 24-inch LED screen (resolution: 1366 × 768 px; refresh rate: 60 Hz) with their chin and forehead placed in a height-adjustable chinrest. Stimulus presentation was implemented using Psychophysics toolbox (Version 3.0.14) for MATLAB (R2017a; MathWorks, Natick, MA). They were instructed to move as little as possible and fixate a white cross on a grey background presented on the screen. Spontaneous eye blink rate was recorded using a standard HD webcam placed above the middle of the screen. A single recording duration of 5 minutes has been shown to suffice for assessing stable spontaneous eye blink rate values (Barbato et al., 2000; Zaman & Doughty, 1997). Pupillometry data were collected using a RED-500 remote eye-tracking system (sampling rate (SR): 500 Hz; Sensomotoric Instruments, Teltow, Germany). heart rate data was acquired utilizing BIOPAC Systems hard-and software (SR: 2000Hz; MP 160; Biopac systems, Inc). For cardiovascular recordings an ECG100C amplifier module with a gain of 2000, normal mode, 35 Hz low pass notch filter and 0.5 Hz/1.0 Hz high pass filter was included in the recording system. Disposable circular contact electrodes were attached according to the lead-II configuration. Isotonic paste (Biopac Gel 100) was used to ensure optimal signal transmission.

### Physiological data analyses

#### Spontaneous eye blink rate

Spontaneous eye blink rate per minute was quantified by replaying the recorded videos and counting the single blinks manually by two separate evaluators. In case of a discrepancy >= 2 blinks this was repeated. The mean of both final evaluations was used.

### Cardiac data

Heart rate data (5 min recordings) were visually screened and manually corrected for major artifacts. We used custom MATLAB code to detect each R peak within the raw data. Specifically, the data was down sampled from 2000 to 200 Hz. Next, we applied the inverse maximum overlap discrete wavelet transform from MATLAB’s Wavelet Toolbox to reconstruct the heart rate data based on the 2^nd^ to the 4^th^ scale of the respective wavelet coefficients. The reconstructed data was squared and respective peaks were detected via MATLAB’s findpeaks function. The obtained peak locations were then subjected to a custom R script that used the RHRV toolbox for R (Rodríguez-Liñares et al., 2011) to compute heart rate and standard deviation of normal to normal beat intervals (SDNN) as a heart rate variability measure.

### Pupil data

We used custom MATLAB code for pupil data analysis (5min recordings). First, NaN values as a result of eye blinks or the like were removed. Next, pupil data were down sampled from 500 Hz to 50 Hz. Outliers within a moving window of five seconds (250 data points) (mean ± 2 SD) were removed and linearly interpolated. The remaining data was smoothed using a robust weighted least squares local regression (‘rloess’ function within MATLAB’s Curve Fitting toolbox) with a span of 10 data points. Mean pupil dilation size and its coefficient of variation (standard deviation/mean*100; Aminihajibashi et al., 2019) were then computed and averaged over both hemispheres for each assessment.

### Tyrosine/ placebo administration

Following previous tyrosine administration studies (Hase et al., 2015; Jongkees et al., 2015), participants were administered a dosage of 2 g tyrosine (supplied by The Hut.com Ltd.) or placebo (2 g cellulose powder (Sigma-Aldrich Co.), solved in 200 ml orange juice (Solevita, Lidl). According to Tam et al. (1990), tyrosine administration enhances tyrosine hydroxylation and causes plasma tyrosine levels to peak approximately 60-120 minutes following intake and remain significantly boosted for up to 8h (Glaeser et al., 1979). Consistent with this, the delay between tyrosine/placebo administration and start of the cognitive assessment was exactly 75 minutes. To test whether the participants were blind to the supplementation condition, we asked them to guess their treatment assignment (placebo-tyrosine / tyrosine - placebo) subsequent to completing the second session.

### Sequential RL task

Participants performed 300 trials of a modified version of the original two-step task by Daw et al. (2011). Based on suggestions by Kool et al. (2016) we modified the outcome stage by replacing the fluctuating reward probabilities (reward / no reward) with fluctuating reward magnitudes (Gaussian random walks with reflecting boundaries at 0 and 100, and standard deviation of 2.5).

In short, each trial comprised two successive decision stages. In the 1^st^ stage (S1), participants chose between two options represented by abstract geometrical shapes. Each S1 option probabilistically led to one of two 2^nd^ stage (S2) states that again comprised two choice options represented by abstract geometrical shapes. Which S2 stage state was presented depended probabilistically on the S1 choice according to a fixed common (70% of trials) and rare (30% of trials) transition scheme. The S2 stage choice options each led to a reward outcome. To achieve optimal performance, participants had to learn two aspects of the task. They had to learn the transition structure, that is, which S1 stimulus preferentially (70%) led to which pair of S2 stimuli. Further, they had to infer the fluctuating reward magnitudes associated with each S2 stimulus. We used different but matched task versions for the two testing days (tyrosine/placebo, counterbalanced). Task versions used different S1 and S2 stimuli, and different S2 random walks. However, S2 Gaussian random walks were matched on variance and mean across task versions.

On each testing day, participants underwent extensive self-paced, computer-based instructions. Instructions provided detailed information about the task structure, the fixed transition probabilities between stages S1 and S2 and the fluctuating reward outcomes in S2. Participants were instructed to earn as much reward points as possible, and that following task completion reward points were converted to € such that they could win a bonus of up to 4€ in addition to their reimbursement of 10€/h. They performed 20 practice trials prior to task performance, with different random walks and stimuli.

### Temporal Discounting Task

On each testing day, following the sequential RL task, participants performed 128 trials of a temporal discounting task. On each trial, participants selected between a “smaller sooner” (SS) reward available immediately and a “larger later” (LL) reward available after a particular delay in days. SS rewards were fixed to 20 € throughout the trials. LL magnitudes were computed by multiplying the SS reward with [1.01, 1.02, 1.05, 1.10, 1.15, 1.25, 1.35, 1.45, 1.65, 1.85, 2.05, 2.25, 2.65, 3.05, 3.45, 3.85] or 1.01, 1.03, 1.08, 1.12, 1.20, 1.30, 1.40, 1.50, 1.60, 1.80, 2.00, 2.20, 2.60, 3.00, 3.40, 3.8]. The temporal delays of the LL rewards ranged from 1 day to 122 days in eight steps [1, 3, 5, 8, 14, 30, 60,120] days or [2, 4, 6, 9, 15, 32, 58, 119], respectively. LL magnitudes and delays were counterbalanced across the testing days and participants. As in previous studies (Green et al., 1997; Wagner et al., 2020) all choice options were hypothetical. Notably, discount rates for real and hypothetical rewards show a high correlation and similar neural underpinnings (Bickel et al., 2012).

Both task were implemented using the Psychophysics Toolbox Version 3 (PTB-3) running under MATLAB (The MathWorks ©).

### Demographic and psychological screening

At the end of the first testing day, participants completed a brief survey on demographic data, socioeconomic status, weight and height (for calculating Body Mass Index (BMI)), trait impulsivity (Barratt-Impulsiveness Scale (BIS-15; German Version: Meule et al., 2020; Spinella, 2007), Behavioral Inhibition and Behavioral Activation System (BIS/BAS scale; German Version: Strobel et al., 2001; Carver & White, 1994), and severity of depressive symptoms (Beck Depression Inventory-II (BDI-II; German Version: Kühner et al., 2007; Beck et al., 1996). Since depressive symptoms can be linked to altered dopamine transmission (Belujon & Grace, 2017), one participant was excluded from data analysis (BDI-II score > 28). Another participant was excluded due to a diagnosed psychiatric condition in the past.

### Model free analysis

For the sequential RL task we used (generalized) mixed effects regression models to examine S1 stay/shift choice patterns, as well as S1 and S2 RTs. These response variables were modeled as a function of previous reward, state transition (common vs. rare) and supplementation (tyrosine vs. placebo). As a model-agnostic measure of temporal discounting, we examined arcsine-square-root transformed proportions of LL choices as a function supplementation condition (tyrosine vs. placebo) with participants as a random effect using a generalized mixed effects regression approach. In line with our modeling analyses (see below), data were filtered such that implausibly fast RTs (see details on DDM modeling) were excluded.

### Computational modeling and data analysis

#### Sequential RL task

We implemented a modified version of the original RL model by Daw et al. (2011) to analyze model-free and model-based contributions to behavior. The model updates model-free state-action values (*Q*_*MF*_-values, Eq. 7, 8) in both stages S1 & S2 via a prediction error scheme (Eq. 3, 4). In S1, model-based state-action values (*Q*_*MB*_) are then computed from the transition and reward estimates using the Bellman Equation (Eq. 5). To account for potential modulatory effects of tyrosine vs. placebo supplementation, we included additive ‘shift’ parameters *s*_*x*_ for each parameter *x* that were multiplied by dummy-coded supplementation predictors *I*_*t*_ (= 1, *TYR*; = 0, *PLC*).

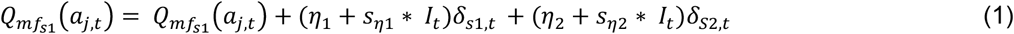

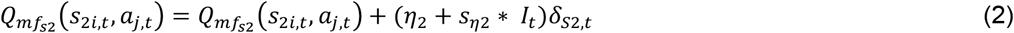

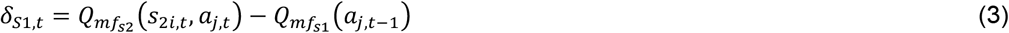

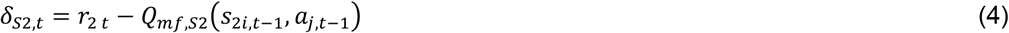

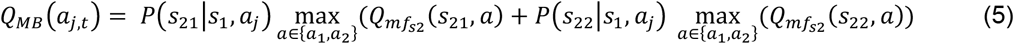

Here, *i* indexes the two different second stage stages (*S*_21_, *S*_22_), *j* indexes actions *a* (*a*_1_, *a*_2_) and *t* indexes the trials. Further, *η*_1_ and *η*_2_denote the learning rate for S1 and S2, respectively. S2 model-free *Q*-values are updated by means of reward (*r*_2,*t*_) prediction errors (*δ*_*S*2,*t*_) (Eq. 8, 10). To model S1 model-free *Q*-values we allow for reward prediction errors at the 2nd-stage to influence 1st-stage *Q*-values (Eq. 2, 4). In addition, *Q*-values of all unchosen stimuli were assumed to decay with decay-rate *γ*_*s*_ (Toyama et al., 2017, 2019) towards the mean of the reward outcomes (0.5) according to Eq. 6 and 7:

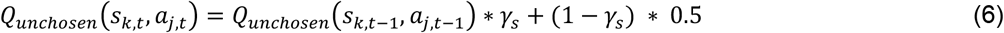

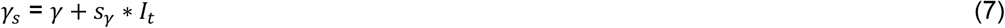

with *k ∈* {1, 21, 22} indexing the first (S1) and the two second stage (S21, S22) stages.

This learning model was then combined with two different choice rules: softmax action selection (see *SI*), and the drift diffusion model (Shahar et al. 2019).

### Drift diffusion model (DDM) implementation

We replaced the standard softmax action selection (see *SI*) with a series of drift diffusion model (DDM)-based choice rules to more comprehensively examine modulatory effects of tyrosine (vs. placebo) supplementation on choice dynamics. The DDM belongs to the family of sequential sampling models. In these models, binary decisions are assumed to arise from a noisy evidence accumulation process that terminates as soon as the evidence exceeds one of two response boundaries. For each stage S1/S2 of the seq. RL task, the upper boundary was defined as selection of one stimulus, whereas the lower boundary was defined as selection of the alternative stimulus. RTs for choices of the alternative option were multiplied by −1 prior to model fitting. Prior to modeling, we filtered the choice data using percentile-based RT cut-offs, such that on a group-level the fastest and slowest one percent of all trials according to RTs were excluded from modeling, and on an individual subject level the fastest and slowest 2.5 percent were further discarded. With this we avoid that outlier trials with implausible short or long RTs bias the results. We then first examined a null model (DDM_0_) without any value modulation. Trial-wise RTs on each stage S1 & S2 are assumed to be distributed according to the Wiener-First-Passage-Time (*wfpt*):

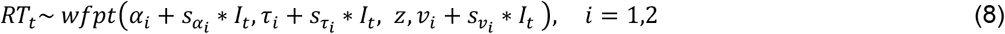

Here, boundary separation parameters *α*_*i*_ model the amount of evidence required before committing to a decision in each stage Si, non-decision time *τ*_*i*_ model components of the RT that are not directly implicated in the choice process, such as motor preparation and stimulus processing. The starting point bias *z* models a bias towards one of the response boundaries before the evidence accumulation process starts. This was set to .5 for both stages, as the boundaries were randomly associated with the choice options on each stage. The drift-rate parameters *v*_*i*_ model the speed of evidence accumulation. Note that for each parameter *x*, we also included a parameter *s*_*x*_ that models potential modulatory effects of tyrosine (vs. placebo) supplementation (coded via the dummy-coded condition predictor *I*_*t*_).

As in previous work (Fontanesi et al., 2019; Pedersen et al., 2017; Peters & D’Esposito, 2020), we then set up hybrid RL DDMs with modulation of drift-rates by value differences between the respective choice options, separately for each stage. First, we set up a model with a linear modulation of drift-rates (DDM_lin_) (Pedersen et al., 2017). For S1, this yields

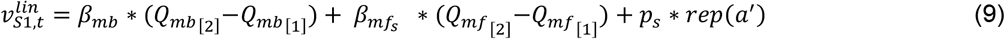

and the drift-rate in S2 is calculated as

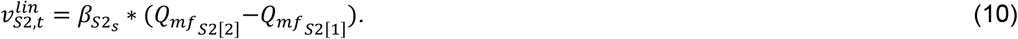

with

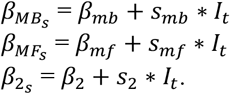

We next set up a DDM with non-linear (sigmoid) drift-rate modulation (DDM_S_) that has recently been shown to better account for the value-dependency of RTs compared with the DDM_lin_ (Bartels et al., 2018; Fontanesi et al., 2019; Peters & D’Esposito, 2020; Wagner et al., 2020, 2021). In this model, the scaled value difference from Eq. 9 & 10 are additionally modulated by a sigmoid function with asymptote

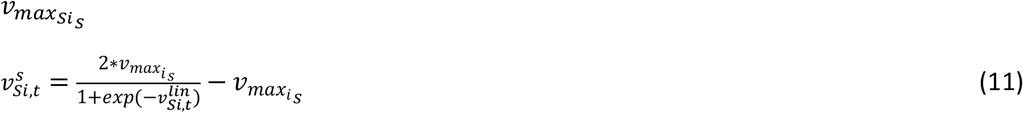

with

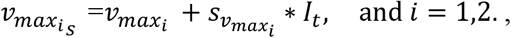

### Temporal discounting model

We applied a single-parameter hyperbolic discounting model to describe how subjective value changes as a function of LL reward magnitude and delay:

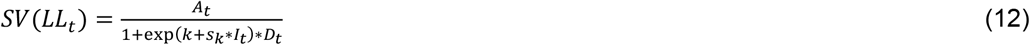

Here, *A*_*t*_ is the reward magnitude of the LL option on trial *t, D*_*t*_ is the LL delay in days on trial *t* and *I*_*t*_ denotes the dummy-coded predictor of the supplementation condition. The model has two free parameters: *k* is the hyperbolic discounting rate (here modeled in log-space) and *s*_*k*_ models a potential additive effect of tyrosine (vs. placebo) on temporal discounting.

### Drift diffusion implementation

In a next step, similar to modeling of the seq. RL data, we replaced softmax action selection (see *SI*) with a series of drift diffusion model (DDM)-based choice rules. In all DDM implementations, the upper boundary was defined as the selection of the LL option, whereas the lower boundary was defined as choosing the SS option. RTs for choices of the SS option were multiplied by −1 prior to model fitting. We used a percentile-based cut-off similar to the one described in the seq. RL modeling section. We again first implemented a null model (DDM_0_) without any value modulation of the drift-rate *v*:

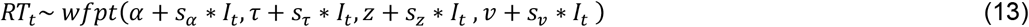

In contrast to the RL model, here the starting point bias *z* was fitted to the data, such that z>.5 reflected a bias towards the LL boundary, and z<.5 reflected a bias towards the SS boundary. As in previous work (Fontanesi et al., 2019; Pedersen et al., 2017; Peters & D’Esposito, 2020a), we then set up temporal discounting DDMs with a modulation of drift-rates by the difference in subjective values between the LL and SS options. First, we set up an implementation with a linear modulation of drift-rates (DDM_lin_) (Pedersen et al., 2017):

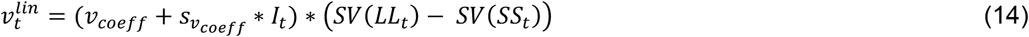

We next examined a DDM with non-linear (sigmoid) trial-wise drift-rate scaling (DDM_S_) that has recently been reported to account for the value-dependency of RTs better than the DDM_lin_ (Fontanesi et al., 2019; Peters & D’Esposito, 2020; Wagner et al., 2020). In this model, the scaled value difference from Eq. 17 is additionally passed through a sigmoid function with asymptote v_max_:

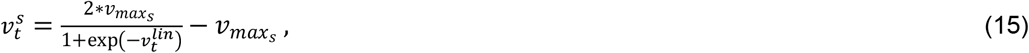

with

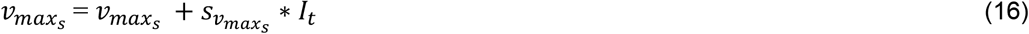

All parameters were again allowed to vary according to the supplementation condition, such that we included *s*_*x*_ parameters for each parameter *x* that were multiplied with the dummy-coded condition predictor *I*_*t*_.

### Hierarchical Bayesian model estimation

Models were fit to all trials (after exclusion of implausibly short or long RTs) from all participants using a hierarchical Bayesian modeling approach with separate group-level distributions for all parameters of the placebo (baseline) condition and additional shift parameters *s*_*x*_ to model tyrosine specific effects on all parameters. Model fitting was performed using MCMC sampling as implemented in STAN (Stan Development Team, 2020) running under R (Version 3.5.1) and the rSTAN package (Version 2.21.0). For baseline group-level means, we used uniform and normal priors defined over numerically plausible parameter ranges (see code and data availability section for details). For all *s*_*x*_ parameters modeling tyrosine-related effects on model parameters, we used normal priors with means of 0 and numerically plausible standard deviations (range=1-10). For group-level standard deviations we used half-cauchy distributed priors with location=0 and scale=2.5 for the temporal discounting data and uniform priors within pausible ranges (mean=0, range standard deviation=10-50). Sampling was performed with four chains, each chain running for 4000 (6000 for the seq. RL data) iterations without thinning after a warmup period of 3000 (5000 for the seq. RL data) iterations. Chain convergence was then assessed via the Gelman-Rubinstein convergence diagnostic 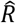 with 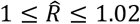 as acceptable values. For both tasks, relative model comparison was performed via the *loo*-package in R (Version 2.4.1) using the Widely-Applicable Information Criterion (WAIC) and the estimated log pointwise predictive density (elpd) which estimates the leave-one-out cross-validation predictive accuracy of the model (Vehtari et al., 2017). We then report posterior group distributions for all parameters of interest as well as their 80% and 90% highest density intervals (HDI). For tyrosine-related effects, we report Bayes Factors for directional effects of parameter distributions of *s*_*x*_, estimated via kernel density estimation using R via the RStudio (Version 1.3) interface. These are computed as the ratio of the integral of the posterior difference distribution from 0 to +∞ vs. the integral from 0 to −∞. Using common criteria (Beard et al., 2016), we considered Bayes Factors between 1 and 3 as anecdotal evidence, Bayes Factors above 3 as moderate evidence and Bayes Factors above 10 as strong evidence. Bayes Factors above 30 and 100 were considered as very strong and extreme evidence respectively, whereas the inverse of these reflect evidence in favor of the opposite hypothesis.

### Posterior Predictive checks

We carried out posterior predictive checks to examine whether models could reproduce key patterns in the data, in particular the value-dependency of RTs(Peters & D’Esposito, 2020) and of participant’s choices. For the seq. RL task, we extracted 500 unique combinations of posterior parameter estimates from the respective posterior distributions and used these to simulate 500 datasets using the Rwiener package (Version 1.3.3). We then show median RTs of observed data and the median RTs from the 500 simulated datasets for all DDMs as a function of value differences. Value differences for S1 were computed as the absolute difference between the maximum 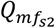 values of each S2 stage weighted by their respective transition probability. Value differences in S2 were computed as the difference in the actual reward values of the respective choice options. Similarly, we show that our models capture the dependency of participants’ stay probabilities in S1 on S1 value differences, and the dependency of their fraction of optimal (max[reward]) choices in S2 on S2 reward differences. For the intertemporal choice task, we binned trials of each individual participant into five bins, according to the absolute difference in subjective LL vs. SS (“decision conflict”), computed according to each participant’s median posterior *k* parameter from the DDM_S_ separately for the placebo and tyrosine condition. For each participant and condition, we then plotted the mean observed RTs and the percentage of LL choices as a function of decision conflict, as well as the mean RTs and fraction of LL choices across 500 data sets simulated from the posterior distributions of the DDM_0_, DDM_lin_ and DDM_S_ and softmax model (choices only).

## Results

### Blindness to the supplementation regime

We assessed whether participants were able to guess on which testing day they received tyrosine. Notably, with 13 (46 %) participants having guessed their supplementation regime correctly we found that this was not above chance level (χ^2^ = .143, p = .706).

### Physiological data

Participants underwent two physiological arousal assessments per session (five minutes each), assessing spontaneous eye blink rate, pupil dilation and heart rate, as well as variability of the latter two. The first assessment was conducted after participants entered the lab as a baseline (t0). The second assessment was conducted 60 minutes following placebo/tyrosine administration (t1). For each measure, we computed the intraclass correlation coefficient to assess the test-retest reliability between the two baseline sessions, and % change at t1 compared to t0 following placebo and tyrosine to assess potential tyrosine-related modulation of physiological arousal, respectively. As expected, spontaneous eye blink rate showed substantial inter-individual variability (range=1.58-48.26; Figure 1a), but proved to be a good test-retest reliability (Table1; Figure 1a). Tyrosine did not significantly modulate spontaneous eye blink rate compared to placebo (Table 1; Figure 1d). Pupil dilation and variability (standard deviation/mean*100) showed moderate and good reliability, respectively (Table 1; Figure 1b). While pupil dilation size was not significantly affected by tyrosine (Table 1), changes in pupil dilation fluctuation between t1 and t0 significantly differed between tyrosine and placebo (Table 1; Figure 1e). and heart rate variability also showed moderate to good test-retest reliability across t0 sessions (Table 1; Figure 1c). Tyrosine compared with placebo was associated with a stronger heart rate deceleration in t1 compared with t0 (Table 1; Figure 1f). There was no significant tyrosine-related modulation of heart rate variability between t1 and t0 (Table 1). Note that in the light of the exploratory nature of the physiological data analyses, we report uncorrected p-values.

**Table 1.** Mean (standard error) of physiological arousal markers during the two baseline sessions, test-retest reliability (intraclass correlation coefficient, p-value), and related percent change thereof 1h post placebo / tyrosine (PLC / TYR) intake. The last column depicts paired t-tests computed on changes following tyrosine vs. placebo. SEBR: spontaneous eye blink rate per minute; PD: mean pupil dilation size; PDV: pupil dilation variability (standard deviation/mean*100); HR: heart rate; HRV: heart rate variability – standard deviation of normal-to-normal heart beat intervals.

| | baseline | test-retest icc (p) | ΔPLC [%] | ΔTYR [%] | $\Delta$(TYR-PLC) t (p) |
| --- | --- | --- | --- | --- | --- |
| SEBR | 16.84 (±2.24) | .84 (4.62*10<sup>-9</sup>) | -5.32 (±5.87) | -2.92 (±7.41) | .31 (.76) |
| PD | 3.55 (±.06) | .59 (3.41*10<sup>-4</sup>) | -1.38 (±1.23) | -.26 (±1.57) | .75 (.46) |
| PDV | 10.21 (±.72) | .89 (4.06*10<sup>-11</sup>) | -6.74 (±2.8) | 8.06 (±5.28) | **3.34 (.002)** |
| HR | 69.98 (±1.96) | .78 (1.87*10<sup>-7</sup>) | -5.02 (±1.33) | -7.72 (±1.18) | **-2.36 (.03)** |
| HRV | 62.03 (±3.76) | .74 (1.87*10<sup>-6</sup>) | 11.94 (±5.4) | 15.86 (±4.39) | .68 (.5) |

**Figure 1.**
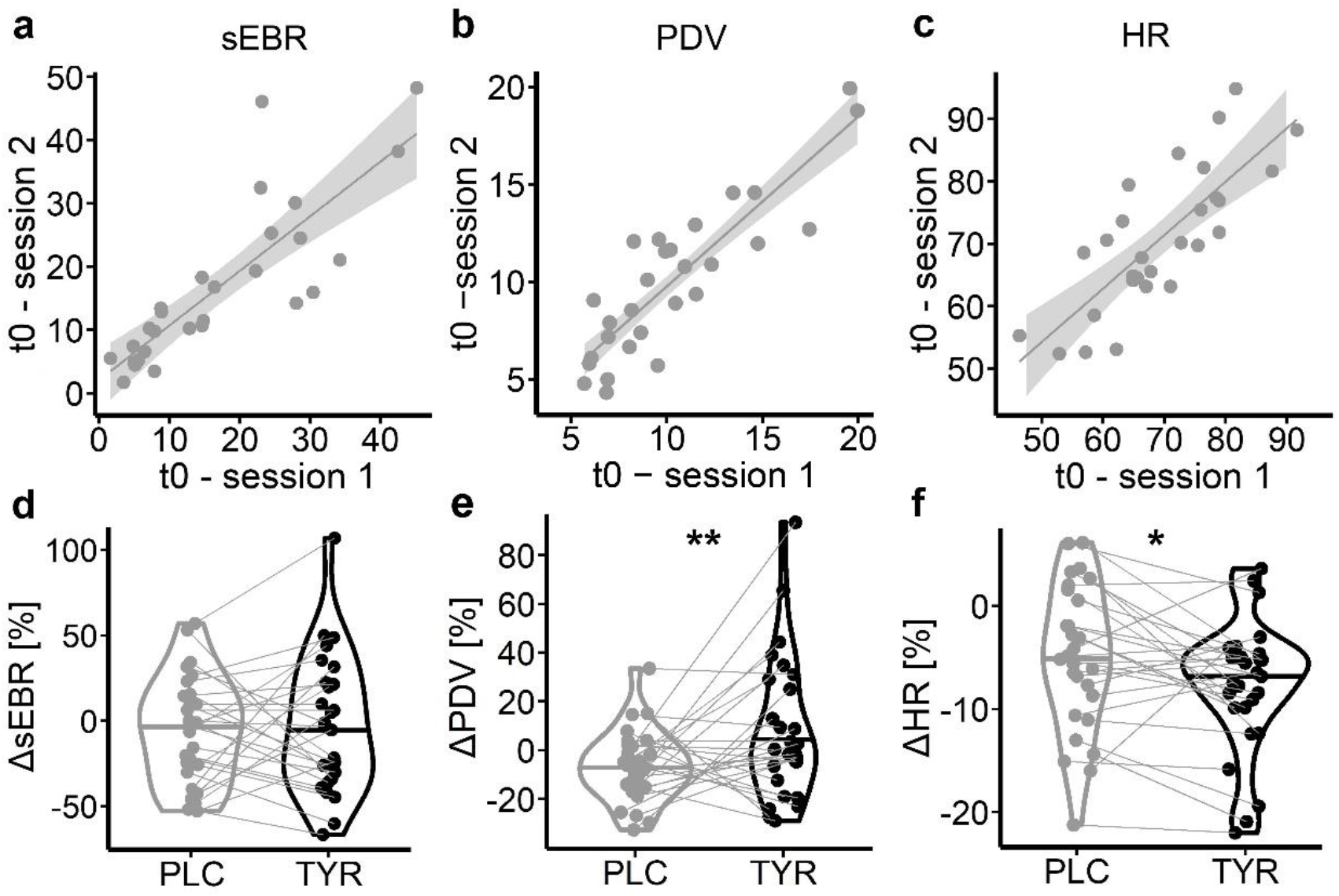
a-c: test-retest reliability across the two baseline (t0) sessions for (a) spontaneous eye blink rate (spontaneous eye blink rate, blinks per minute), (b) pupil dilation (PD) variation (standard deviation/mean*100) and (c) heart rate (HR). d-f: changes per measure between t0 and 60min post placebo or tyrosine supplementation (t1). * denotes a significant (p<.05) difference between tyrosine and placebo associated changes from t0 to t1.

### Sequential RL task

#### Model-agnostic analyses

Tyrosine had no significant effect on participants’ average payout per trial (tyrosine: 61.39 (± .72) vs placebo: 62.25 ± .5 [mean (±SE)]; t(27) = −1.64, p=.11). We used a generalized mixed effects regression approach to examine participants’ stay/shift behavior in S1 as a function of previous reward receipt, state transition (common vs. rare) and supplementation (tyrosine vs placebo) allowing for interactions including participants as a random effect. We observed a main effect of reward (*β*=.13, SE=.04, p=1.09*10^-3; Table 2), reflecting an model-free contribution to behavior, and a reward × transition interaction indicating that subjects incorporated a model-based RL strategy (*β*=-.54, SE=.07, p=1.22*10^-14; Table 2). We observed neither a significant effect of condition (tyrosine vs. placebo) nor a significant interaction effect with condition on stay probability in S1.

**Table 2.** Effects on participants’ probability to choose the same option in S1 as in the previous trial from a generalized mixed effects regression (glmer) analysis (rew=reward; trans=state transition; TYR=tyrosine; glmer model: stay(S1) ∼ rew*trans*TYR + (rew*trans + 1 | participant)). And effects of state transition (trans), TYR vs. placebo (TYR) supplementation and their interaction on participants S2 RTs from a lmer analysis (lmer model: RT(S2)∼ trans*TYR + (trans + 1 | participant).

| S1 – stay probability | $\beta$ | SE | z/t | p |
| --- | --- | --- | --- | --- |
| intercept | 1.4 | .14 | 9.71 | **2.63\*10<sup>-22</sup>** |
| rew | .13 | .04 | 3.27 | **1.09\*10<sup>-3</sup>** |
| trans | -.06 | .03 | -1.71 | .09 |
| TYR | .01 | .02 | .54 | .59 |
| rew*trans | -.54 | .07 | -7.71 | **1.22\*10<sup>-14</sup>** |
| rew*TYR | .03 | .02 | 1.26 | .21 |
| trans*TYR | -.04 | .02 | -1.44 | .15 |
| rew*trans*TYR | .03 | .02 | 1.21 | .23 |
| S2 – RT |  |  |  |  |
| intercept | 0.7 | 0.02 | 45.74 | **3.97<sup>-27</sup>** |
| trans | .06 | 7.2*10<sup>-3</sup> | 8.76 | **2.28\*10<sup>-9</sup>** |
| TYR | -4.91*10<sup>-3</sup> | 1.83*10<sup>-3</sup> | -2.67 | **7.63\*10<sup>-3</sup>** |
| trans*TYR | 5.08*10<sup>-3</sup> | 1.84*10<sup>-3</sup> | 2.76 | **5.82\*10<sup>-3</sup>** |

In a similar fashion we examined possible effects of tyrosine supplementation on participants’ S1 and S2 RTs. We first tested for a general effect of tyrosine supplementation on trial-wise RTs in a mixed effects regression model, including supplementation as a fixed effect and transition and subjects as random effects. Participants responded faster under tyrosine compared to placebo (*β* =-6.61*10^-3, SE=1.19*10^-3, p=2.63*10^-8, Figure 2a). Increased S2 RTs following rare transitions is another indicator of model-based control similar to the above reported reward*transition interaction in S1 choice probabilities (Otto et al., 2015; Shahar et al., 2019). Thus, next we computed a mixed effects regression with transition, supplementation and their interaction as fixed effects on S2 RTs, and subject as a random effect. As previously reported (Otto et al., 2015; Shahar et al., 2019), we observed a significant main effect of transition on S2 RTs (*β*=.06,SE=7.2*10^-3, p=2.28*10^-9), Figure 2b).

**Figure 2.**
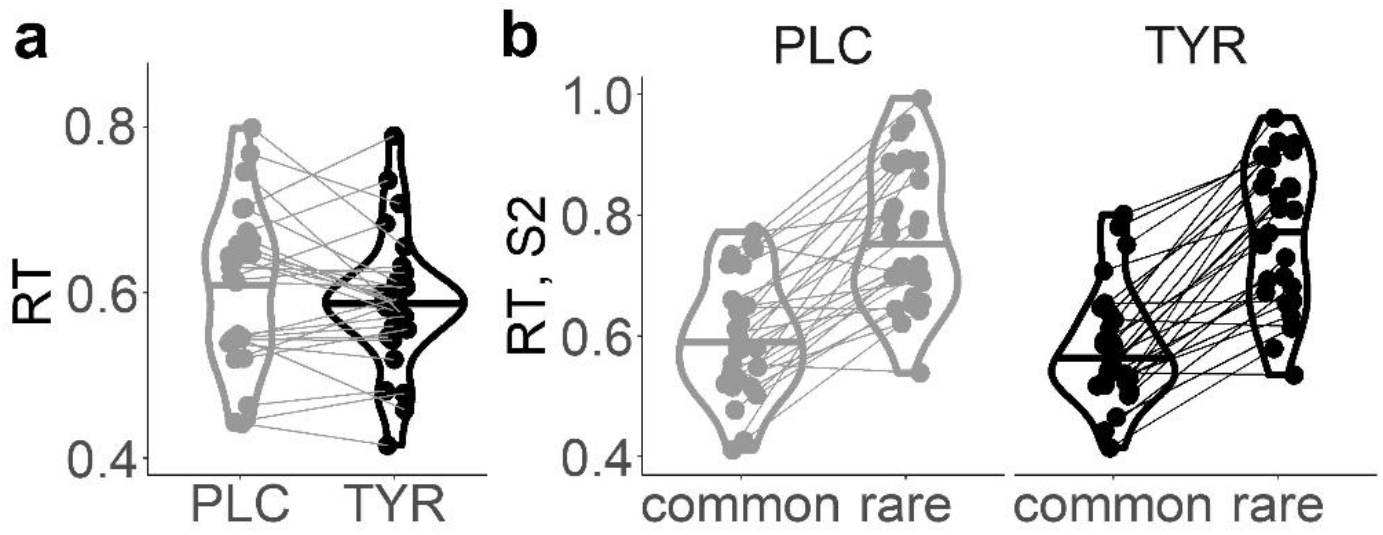
Violin plots of (a) median response times (RT) over both stages S1 & S2, and of (b) RTs in S2 subsequent to a common vs. a rare state transition from the sequential RL task per supplementation condition (placebo vs. tyrosine).

Tyrosine significantly reduced S2 RTs (*β*=-4.91*10^-3, SE=1.83*10^-3, p=7.63*10^-3, Table 1 b). Furthermore, a significant transition*tyrosine interaction (*β*=5.08*10^-3.06, SE=1.84*10^-3, p=5.82*10^-3, Table 1 b, Figure 1b) reflected a more pronounced slowing of RTs following rare vs. common transitions under tyrosine compared to placebo. Taken together, RT analyses suggest a general RT reduction and an increase of model-based RL following tyrosine. Notably, in our modified task version and in contrast to the original sequential RL task from Daw et al., (2011), increased model-based RL usually leads to increased payout (Kool et al., 2016). This was reflected in a significant association of participants’ mean payoffs per trial and S2 RT differences between rare and common transitions (placebo(tyrosine): r=.63(.64), p=3.45*10^-4(2.47*10^-4); Figure 3)

**Figure 3.**
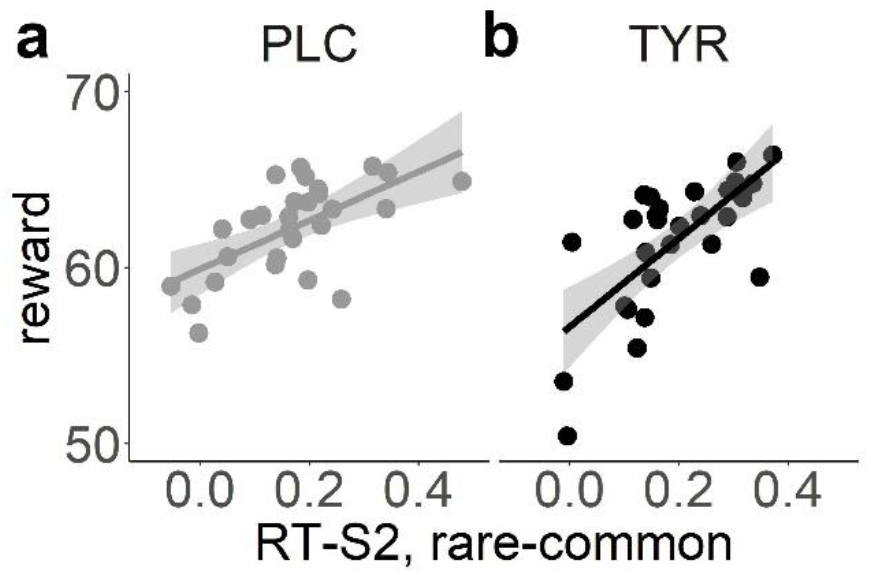
Correlation of participants’ mean rewards per trial and the differences in S2 RTs following a rare vs. a common state transition for the placebo (left) and tyrosine (right) condition.

### Sequential RL task, DDM

We used a DDM choice rule to make use of both choices and RT distributions (Shahar et al., 2019, Wagner et al., 2021). First, we examined the model fit (WAIC, elpd) of three implementations of the DDM that varied in the way they accounted for the modulation of trial-wise S1 & S2 drift-rates by Q-value differences. The DDM with nonlinear drift-rate scaling (DDMs) (Fontanesi et al., 2019; Peters & D’Esposito, 2020; Wagner et al., 2020) accounted for the data best when compared to a DDM with linear scaling (DDM_lin_) (Pedersen et al., 2017) and a null model without value modulation (DDM_0_) (see *SI*, Table S3). Note, that we also compared the three DDMs with a softmax model formulation according to the proportion of correctly predicted binary choices in S1 and S2, respectively. Since the softmax model only accounts for choice data, it shows the highest predictive accuracy. However, the DDM_s_ predicted participants’ choices better than the DDM_lin_ and the DDM_0_ (see *SI*, Table S4). Posterior predictive checks for the winning model DDM_s_ showed that it reproduced the effect decision conflict on participants’ RTs and choice patterns (see *SI*, Figure S1).

We confirmed the expected positive associations between trial-wise drift rates (i.e. *β* weights for model-free-, model-based-, and S2 stage Q-values) and Q-value differences (*see SI*, Figure S5 e-g, all 90% HDIs > 0). Model-free and model-based *β* weights were also comparable to the parameters observed in the softmax model (*see SI*, Figure S4 i, j), indicating robust model-free and model-based contributions during RL. Tyrosine administration increased model-based control and choice perseveration (90% HDI > 0; Figure 4 f,m; Table 3) in stage S1. Likewise, the S1 drift-rate asymptote *v*_*max*_, i.e. the maximum rate of evidence accumulation, was reduced following tyrosine compared with placebo (90% HDI > 0; Figure 4 h; Table 3). There was also strong evidence for an increase in learning rates in S1 following tyrosine (Figure 4 j; Table 3). In line with the observed reduction in RTs following tyrosine, we found attenuated decision-thresholds (90% HDI < 0; Figure 4 f; Table 3) and a potential moderate reduction (according to dBF; Table 3) in S2 non-decision times in S2 (Figure 4 d; Table 3) under tyrosine. The S2 drift rate coefficient *β*_2_ was also reduced (Figure 4 g, Table 3).

**Table 3.**
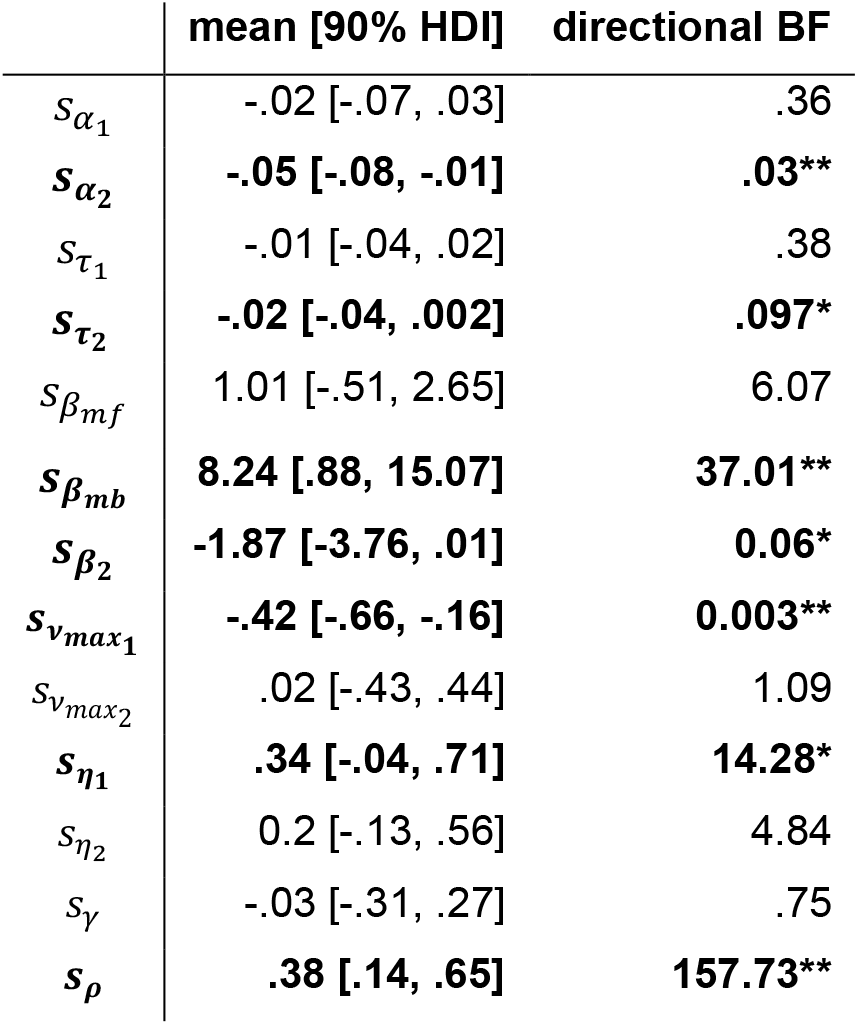
Tyrosine related changes in group-level means of the DDM_s_ for the seq. RL data. We report mean (and 90% HDIs) for the ‘shift’ parameters modeling tyrosine-related effects, and Bayes Factors for directional effects (tyrosine>placebo). * (**): Strong evidence (dBF>10 /<.1) for a tyrosine-related effect (& 90% HDIs outside of zero).

|  | mean [90% HDI] | directional BF |
| --- | --- | --- |
| $s_{\alpha_1}$ | -.02 [-.07, .03] | .36 |
| $s_{\alpha_2}$ | **-.05 [-.08, -.01]** | **.03\*\*** |
| $s_{\tau_1}$ | -.01 [-.04, .02] | .38 |
| $s_{\tau_2}$ | **-.02 [-.04, .002]** | **.097\*** |
| $s_{\beta_{mf}}$ | 1.01 [-.51, 2.65] | 6.07 |
| $s_{\beta_{mb}}$ | **8.24 [.88, 15.07]** | **37.01\*\*** |
| $s_{\beta_2}$ | **-1.87 [-3.76, .01]** | **0.06\*** |
| $s_{v_{max1}}$ | **-.42 [-.66, -.16]** | **0.003\*\*** |
| $s_{v_{max2}}$ | .02 [-.43, .44] | 1.09 |
| $s_{\eta_1}$ | **.34 [-.04, .71]** | **14.28\*** |
| $s_{\eta_2}$ | 0.2 [-.13, .56] | 4.84 |
| $s_{\gamma}$ | -.03 [-.31, .27] | .75 |
| $s_{\rho}$ | **.38 [.14, .65]** | **157.73\*\*** |

**Figure 4.**
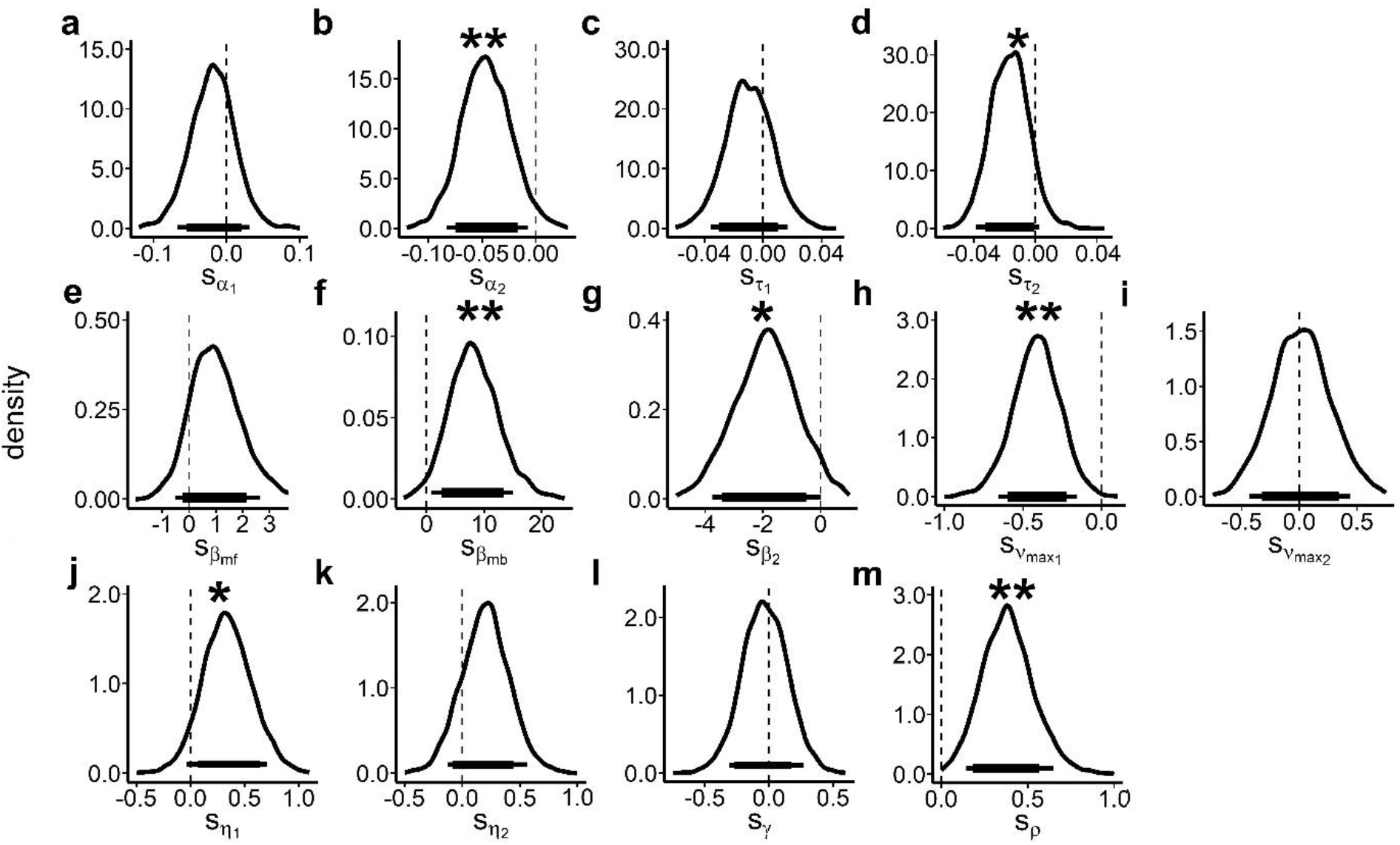
Group-level mean posterior distributions of the tyrosine-related parameter changes (‘shift parameter’ *s*_*x*_) of the DDM_s_ model for the seq. RL task data. Horizontal (thick) lines show 90 (80) % HDIs. Depicted parameters are (a), (b) shift in decision-thresholds 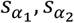 (c), (d) non-decision times 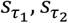; (e)-(g) weights for model-free Q-values 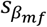, model-based Q-values 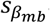 and S2 stage Q-values 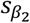; (h), (i) the asymptote of respective drift-rates 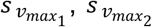; (j), (k) learning-rates in S1 and S2 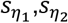, (l) decay rate of unchosen options *s*_*γ*_ and (m) choice perseveration *s*_*ρ*_. * Denote strong evidence for tyrosine-related effects according to directional BF (>10, <.1; see Table 3), and ** denote strong effects (dBF>10, <.1) with 90% HDI outside of zero (Table 3).

### Temporal discounting task

#### Model agnostic analyses

We used a generalized mixed effects regression to test for tyrosine-related effects on trial-wise choices in the temporal discounting task with supplementation condition (tyrosine vs. placebo), larger-but-later (LL) reward magnitude (median split), delay (median split) and their interactions as fixed effects and participants as random effect. The likelihood of choosing the LL option was significantly modulated by delay (*β* =-.95, SE=0.04, p=1.07*10^-11, Table 4), LL magnitude (*β* =1.82, SE=.16, p=3.97*10^-29, Table 4), and their interaction (*β* =-.17, SE=.08, p=.04, Table 4). While we found no main effect of tyrosine on choices (Figure 5 a, Table 4), we observed a significant delay*tyrosine interaction (*β* =.09, SE=.04, p=.02, Table 4) reflecting that following tyrosine participants discounted LL rewards less steeply compared to placebo. Similar to the findings in the seq. RL task, trial-wise RTs were significantly reduced following tyrosine compared to placebo according to a mixed effects regression model including LL magnitude (median split), delay (median split) and supplementation condition and their interaction as fixed effects and participants as random effects (*β* =-.06, SE=6.1*10^-3, p=3.8*10^-23, Table 4, Figure 5 b; see Table 4 for effects of delay & LL magnitude on participants’ RTs).

**Table 4.** Effects on the probability to choose the larger-but-later (LL) option in the temporal discounting task from a generalized mixed effects regression analysis (glmer model: choice(LL)∼delay*LL magnitude*tyrosine + (delay*LL magnitude + 1 | participant)). And effects on participants’ RTs in the temporal discounting task following a lmer model (lmer model: rt ∼ delay*LL magnitude*tyrosine + (delay*LL magnitude +1 | participant)), with delay & LL magnitude as binary predictors via median split in both mixed effects models.

| LL choices | $\beta$ | SE | z | p |
| --- | --- | --- | --- | --- |
| intercept | 1.09 | .44 | 2.5 | .01 |
| delay | -.95 | .04 | -6.8 | **1.07\*10<sup>-11</sup>** |
| LL magnitude | 1.82 | .16 | 11.21 | **3.97\*10<sup>-29</sup>** |
| TYR | .03 | .04 | .66 | .51 |
| delay*LL magnitude | -.17 | .08 | -1.98 | .04 |
| delay*TYR | .09 | .04 | 2.29 | .02 |
| LL magnitude*TYR | -.03 | .04 | -.72 | .47 |
| delay*LL magnitude*TYR | .04 | .04 | 1.18 | .24 |
| RTs |  |  |  |  |
| intercept | 1.23 | .07 | 18.34 | **3.89\*10<sup>-17</sup>** |
| delay | .03 | .01 | 2.46 | .02 |
| LL magnitude | -.04 | .02 | -1.93 | .06 |
| TYR | -.06 | 6.1*10<sup>-3</sup> | -9.95 | **3.8\*10<sup>-23</sup>** |
| delay*LL magnitude | .02 | 8.7*10<sup>-3</sup> | 2.23 | .03 |
| delay*TYR | -3.19*10<sup>-3</sup> | 6.1*10<sup>-3</sup> | -.52 | .6 |
| LL magnitude*TYR | 7.27*10<sup>-3</sup> | 6.1*10<sup>-3</sup> | 1.19 | .23 |
| delay*LL magnitude *TYR | 4.06*10<sup>-3</sup> | 6.09*10<sup>-3</sup> | .67 | .51 |

**Figure 5.**
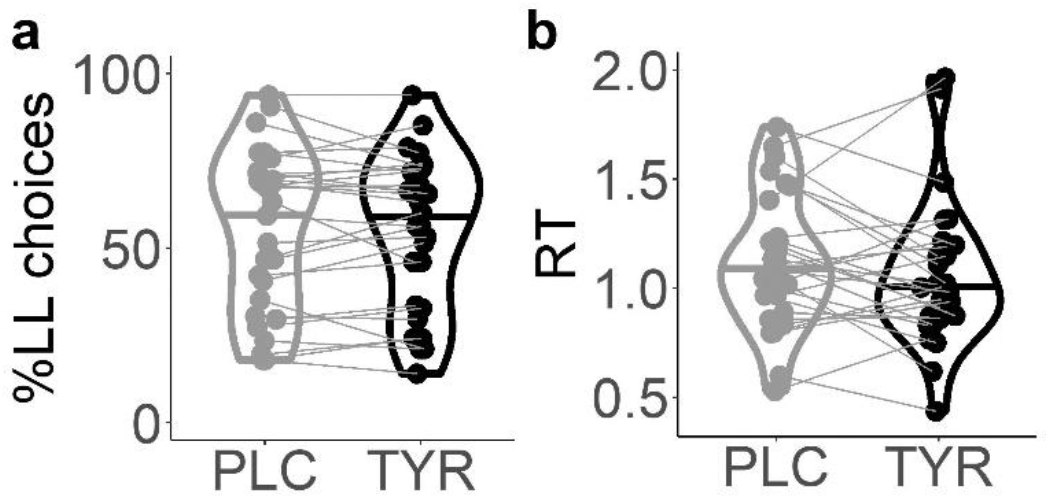
Violin plots of (a) proportions of LL choices following placebo and tyrosine supplementation, and of (b) median response times (RT) in the temporal discounting task per supplementation condition (placebo vs. tyrosine).

### Temporal discounting task, DDM

As for the the seq. RL task data, we first examined the model fit (WAIC, elpd) of three implementations of the DDM that varied in the way they accounted for the modulation of trial-wise drift-rates by value differences. A DDM with nonlinear drift-rate scaling (DDMs) (Fontanesi et al., 2019; Peters & D’Esposito, 2020; Wagner et al., 2020) also accounted for the temporal discounting data best when compared to a DDM with linear scaling (DDM_lin_) (Pedersen et al., 2017) and a null model without value modulation (DDM_0_) (Table 5). We also compared the three DDMs to a softmax model with respect to the proportion of correctly predicted binary choices (SS vs. LL choices). The DDM_s_ predicted participants’ choices numerically on par with the softmax model, whereas the DDM_lin_ and even more so the DDM_0_ performed substantially worse (see *SI Table S3*). Posterior predictive checks of the best-fitting model DDM_s_ revealed that it accurately reproduced the effect of value differences (i.e. decision conflict) on participants’ RTs and the proportion of LL choices (see Supplemental Figure S2). Note that we previously reported extensive parameter recovery analyses for this model (Peters & D’Esposito, 2020; Wagner et al., 2020).

We then examined participants’ discounting behavior in greater detail via the posterior distributions of the group-level mean DDM_s_ parameters (Figure 6). We found no evidence for a general bias towards the SS or the LL option, as the 90% HDI for the starting-point bias parameter *z* overlapped with 0.5 (Figure 6 b). Furthermore, trial-wise drift rates increased with increasing value differences, such that the 90% HDI for the drift-rate coefficient parameter *v* did not include 0 (see Figure 6 h).

**Figure 6.**
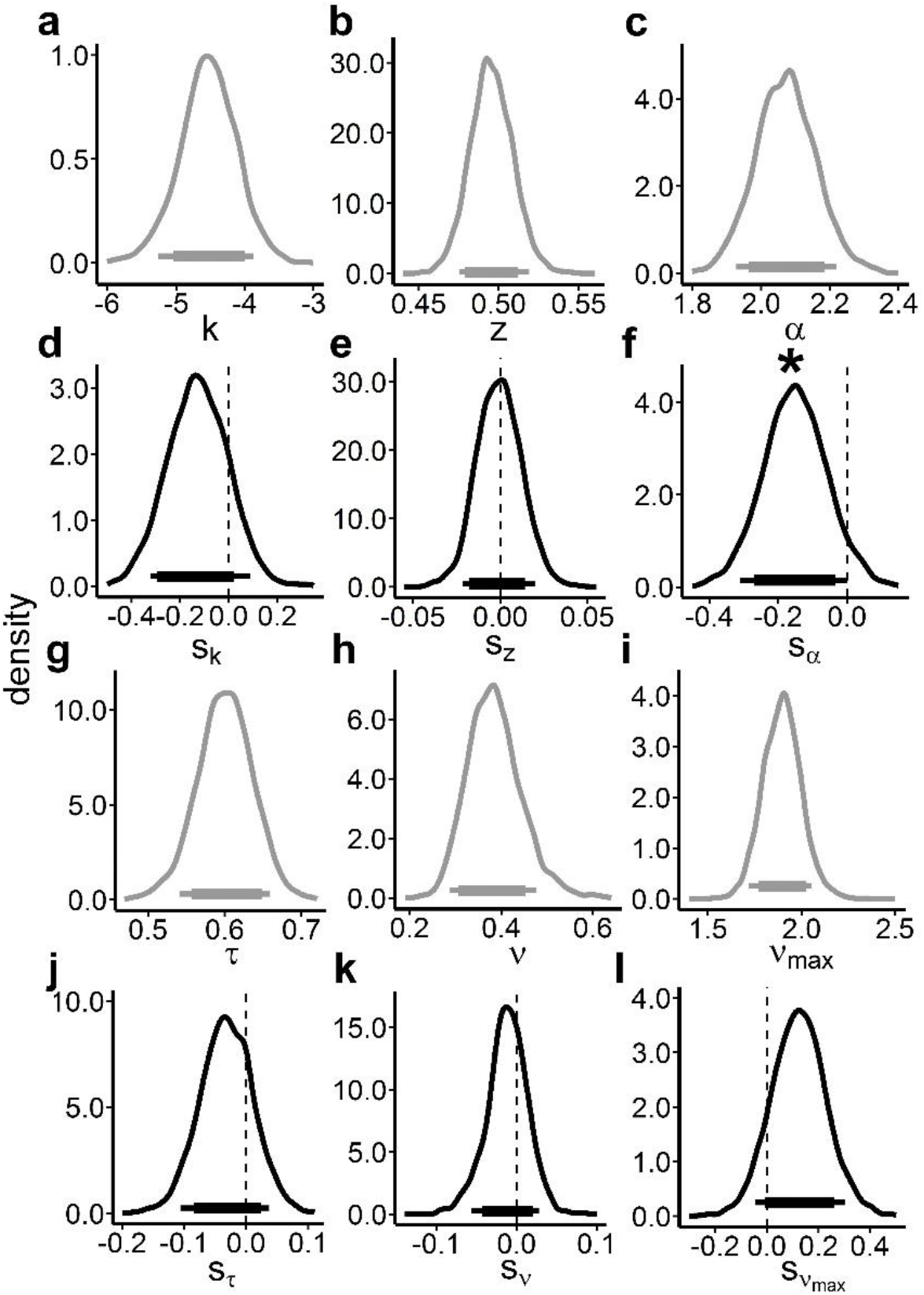
Posterior distributions of the group-level means of the DDM_s_ for the temporal discounting task data for the placebo condition (a-c, g-i; grey plots) and tyrosine-related changes thereof (d-f, j-l; black plots). Horizontal (thick) lines denote 90 (80) % HDIs. Depicted parameters are (a)-(c) discount rate log(*k*), starting point bias *z*, decision-threshold *α*, and (g)-(i) non-decision time *τ*, drift-rate *v* and drift-rate asymptote *v*_*max*_; as well as their tyrosine-related, shifts’ in (d)-(f) & (j)-(l). * Denotes strong evidence for tyrosine-related effects according to directional BF (>10, <.1; see Table 6).

Tyrosine led to a moderate numerical decrease in temporal discounting according to the DDM_s_ (*s*_*k*_; Figure 6 d & Table 6). Furthermore, mirroring the reduced RTs during temporal discounting and similar to the tyrosine-related findings in the seq. RL data, tyrosine substantially reduced decision thresholds (*s*_*α*_; Figure 6 f, Table 6). Furthermore, tyrosine moderately increased the maximum drift rate (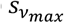; Figure 6 l, Table 6).

**Table 6.** Tyrosine related DDM_s_ for the temporal discounting task data. We report mean and 90% HDIs for the ‘shift’ parameters and Bayes Factors for directional effects (tyrosine>placebo). * denotes strong evidence (dBF>10 /<.1) for a tyrosine-related effect on the respective shift parameter *s*_*α*_.

|  | mean [90% HDI] | directional BF |
| --- | --- | --- |
| $\log(s_k)$ | -.13 [-.32, .09] | .19 |
| $s_z$ | -.00 [-.02, .02] | 0.91 |
| **$s_\alpha$** | **-.15 [-.31, 0]** | **.06\*** |
| $s_\tau$ | -.01 [-.11, .04] | .3 |
| $s_v$ | -0.012 [-.06, .03] | 0.48 |
| $s_{v_{max}}$ | .12 [-.05, .3] | 7.15 |

### Correspondence between task performance and physiology

In exploratory analyses, we tested for associations between task performance (seq. RL and temporal discounting) under placebo and physiological arousal measures (spontaneous eye blink rate, pupil dilation, pupil dilation variability, heart rate and heart rate variability), as well as tyrosine-related changes of these. We restricted these exploratory analyses to core aspects of seq. RL and temporal discounting, and related evidence accumulation processes to reduce multiple testing burden (see *SI* for a detailed description). We found that higher pupil dilation at baseline was associated with more impatient (fewer LL) choices during temporal discounting (% LL choices; r=-.55, p=.002; Figure 7 a). In line, higher pupil dilation was associated with a significant bias towards impatient SS choices (*z*; r=-.63, p=3.0*10^-4; Figure 7 b). With respect to tyrosine-related changes in physiological arousal, greater pre-post heart rate changes following tyrosine compared with placebo were associated with tyrosine-related shifts in temporal discounting (*s*_log(*k*)_; r=.61, p=5.76*10^-4; Figure 7 c). All other tested associations were non-significant after adjusting for False Discovery Rate (Benjamini & Hochberg, 1995; all p-values >=.005; FDR adjusted p-value = .003).

**Figure 7.**
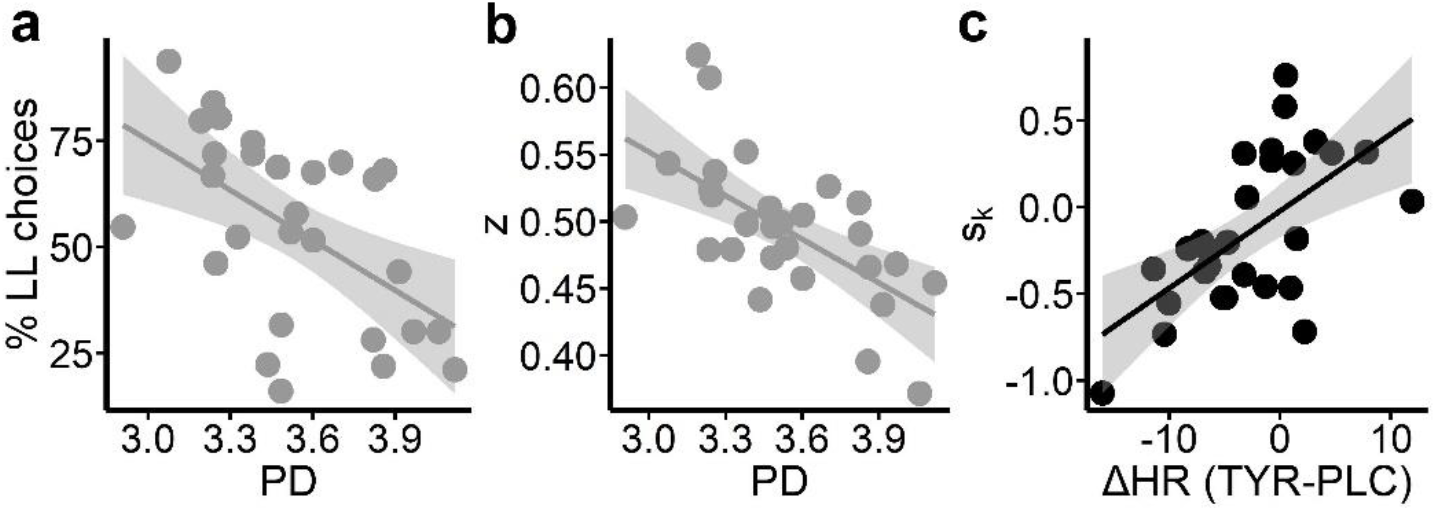
Participants pupil dilation (PD) at baseline was predictive of (a) % LL choices, (b) bias towards SS choices (*z*) and (c) tyrosine associated changes in temporal discounting (*s*_log(*k*)_) was linked with supplementation related differences (tyrosine-placebo) in heart rate (HR) changes relative to baseline.

## Discussion

In a double-blind, placebo-controlled within-subjects design, we investigated the effect of a single dose (2g) of tyrosine supplementation on two fundamental catecholamine dependent decision-making processes-sequential reinforcement learning (seq. RL) and temporal discounting. We used drift diffusion modeling (DDM) in a hierarchical Bayesian approach to reliably quantify tyrosine-related changes within distinct aspects of the dynamic choice processes in both tasks. Assessment of spontaneous eye-blink rate, pupil dilation, heart rate and variability of the latter two both pre and post tyrosine/placebo administration enabled us to associate catecholamine dependent behavior with physiological arousal markers and tyrosine-related modulation of both in an exploratory manner. All physiological measures exhibited moderate to good test-retest reliability. Tyrosine reduced physiological arousal compared to placebo as indexed by modulations of pupil dilation variation and heart rate. On the behavioral level, tyrosine consistently reduced RTs across tasks, without compromising task performance. Hierarchical Bayesian DDMs linked these effects to attenuated decision-thresholds in both tasks. Tyrosine augmented model-based control during seq. RL and, if anything, reduced rather than increased temporal discounting. Participants’ mean pupil dilation at baseline predicted core aspects of their temporal discounting behavior. Tyrosine-related reductions of arousal were associated with individual changes in temporal discounting following tyrosine supplementation.

### Tyrosine attenuates physiological arousal

We assessed pupil dilation (mean & variability), heart rate (mean & variability), as well as spontaneous eye blink rate during a five minute interval at baseline (t0) and 60 minutes following tyrosine/placebo supplementation (t1) on both testing days. First, all three measures exhibited moderate to good test-retest reliability across assessments. T0 assessment was conducted only a few minutes after participants entered the lab. Individual arousal levels were therefore presumably higher at baseline than at t1 (following tyrosine/placebo intake). This was mirrored in a reduced heart rate at t1. Second, the reduction in physiological arousal from t0 to t1 was more pronounced following tyrosine supplementation. This was reflected in a significant difference in the t1-t0 change in pupil dilation and heart rate following tyrosine compared to placebo supplementation. As a precursor of dopamine and noradrenaline, tyrosine supplementation significantly elevates the level of homovanillic acid in rodents and humans, the major catecholamine metabolite, potentially indicating increased catecholamine synthesis and release within the brain (Brodnik et al., 2012; Glaeser et al., 1979; Growdon et al., 1982). In line with this, increased noradrenaline plasma levels have been linked to tyrosine administration in humans (Kishore et al., 2013). Tyrosine can have a hypotensive effect in humans specifically in stressful situations (Deijen et al., 1999). Similar effects in rodents are assumed to be related to *α*-adrenergic receptor stimulation through elevated norepinephrine release (Sved et al., 1979; Yamori et al., 1980). In line, a recent study that tested cognitive and physiological effects of increased norepinephrine transmission reported a stronger heart rate reduction compared to baseline assessment following administration of the norepinephrine antagonist Yohimbine (Herman et al., 2019). These considerations suggest that tyrosine might have attenuated physiological arousal via a norepinephrine-related mechanism.

### Tyrosine reduces RTs

Tyrosine supplementation was consistently associated with reduced RTs, both in temporal discounting and seq. RL. This complements earlier work from Magill et al. (2003) who found that following sleep deprivation, tyrosine significantly reduced RTs compared with placebo across several cognitive tasks. Likewise, acute tyrosine and phenylalanine depletion significantly slowed RTs during a rapid visual information processing task (Roiser et al., 2005). Changes in dopamine and/or norepinephrine neurotransmission might underlie these effects. Recently, Wagner et al. (2020) reported reduced non-decision times following administration of the D2 receptor antagonist Haloperidol in healthy volunteers during a temporal discounting task. They interpreted their finding as facilitated motor responding due to increased dopamine transmission via presynaptic action of low dosages of Haloperidol. Findings of a dopaminergic speeding of RTs in tasks testing response vigor (Guitart-Masip et al., 2011; Hofmans et al., 2021) further support the idea that tyrosine might have reduced RTs via a dopamine-dependent mechanism. However, increased norepinephrine transmission following tyrosine may also have contributed to RT reductions. Pharmacological blockade of central *α*_2_-adrenergic receptors was shown to attenuate RTs, but also increase impulsive responding and increase heart rate and blood pressure in healthy human volunteers (Swann et al., 2013) which however contrasts with our findings regarding temporal discounting performance and physiological correlates of tyrosine intake. However, Herman et al., (2019) found reduced stop-signal RTs indicating attenuated motor impulsivity and stronger heart rate deceleration compared to baseline following a noradrenergic antagonist in comparison to placebo. These inconsistencies may be attributed to the complexity of the noradrenergic system in relation to its cognitive and physiological output depending on the specific receptor family (presynaptic *α*_2_ and postsynaptic *β* and *α*_1_), the major release site and the balance between tonic and phasic transmission (Holland et al., 2021).

### Drift diffusion modeling (DDM)

We implemented value-based decision (temporal discounting) and RL models (model-based RL) in a sequential sampling framework (DDM) (Bruder et al., 2021; Fontanesi et al., 2019; Pedersen et al., 2017; Peters & D’Esposito, 2020b; Shahar et al., 2019; Wagner et al., 2020). Model comparison replicated previous results (Bruder et al., 2021; Fontanesi et al., 2019; Peters and D’Esposito, 2020; Wagner et al. 2020), such that data in both tasks were consistently better accounted for by models assuming a non-linear trial-wise drift rate modulation. Posterior predictive checks of choices and RTs confirmed the superior reproduction of relevant inherent patterns (see *SI*). Prior parameter recovery work confirmed that DDM group-level parameters for temporal discounting data (Peters and D’Esposito, 2020; Wagner et al., 2020) and seq. RL data (Fontanesi et al., 2019; Shahar et al., 2019) recovered well. Notably, this modeling approach allowed us link the tyrosine-related RT reductions in both tasks to a reduction in decision thresholds (see next section). This highlights the valuable insights that comprehensive modeling approaches can add beyond mere analysis of choices and RTs in neuro-cognitive research.

### Tyrosine reduces decision-thresholds

DDM linked the attenuated RTs following tyrosine supplementation to reduced decision thresholds in both tasks. This parameter models the amount of evidence that is accumulated before committing a decision, i.e. the speed-accuracy trade-off (Ratcliff & McKoon, 2008; Smith & Ratcliff, 2004). Interestingly, in neither task was this associated with objective performance decrements: Tyrosine did not reduce participants’ earnings in the seq. RL task nor did it increase impulsive choices during temporal discounting. Conversely, model-based control increased during seq. RL as evident in a tyrosine-related increase in S2 RT slowing following rare transitions and increased *β*_*mb*_ according to the DDM. Likewise, if anything, participants showed a moderate decrease in temporal discounting. Decision thresholds have been linked to subthalamic nucleus activity (Herz et al., 2016). This part of the hyper-direct cortico-basal ganglia pathway (Nambu et al., 2002) receives direct dopaminergic (Canteras et al., 1990; Cragg et al., 2004) and noradrenergic (Canteras et al., 1990; Parent & Hazrati, 1995) input. A further function of the subthalamic nucleus is the withholding of premature responding to allow for more time to evaluate information and select the most appropriate action (Aron et al., 2016; Wessel et al., 2016). Deep brain stimulation of the STN in Parkinsonism was shown to attenuate RTs in a RL task in high conflict trials while leaving RL performance intact (Frank et al., 2007). Tyrosine may have modulated STN activity in a similar manner via increased tonic dopaminergic and or noradrenergic input. We also observed consistent numerical reductions in non-decision times across tasks, although strong evidence was restricted to the second stage in the seq. RL task. This may reflect enhanced motor responding e.g. via increased DA transmission (Wagner et al. 2020) within striato-cortical pathways following tyrosine intake.

### Tyrosine increases model-based control and perseveration in seq. RL

Tyrosine supplementation was associated with increased model-based control during RL as evident in a tyrosine-related increase in S2 RTs slowing following rare transitions and increased *β*_*mb*_ according to the DDM. This convergence across model-agnostic and model-based analyses strengthens the confidence in the result. Although we can only speculate about the potential underlying mechanism, this is in line with previous reports on dopamine precursor L-dopa-related increases in model-based control in healthy participants and Parkinson’s disease patients (Sharp et al., 2016;

Wunderlich et al., 2012). But see Kroemer et al., (2019) for contrary findings. Similarly, tyrosine may have augmented dopamine transmission in fronto-striatal circuits, possibly facilitating model-based prediction error encoding (Daw et al., 2011; Deserno, 2015; Doll et al., 2012). Tyrosine may also have increased dopaminergic and/or noradrenergic input to the hippocampus, potentially facilitating model-based RL (Vikbladh et al., 2019).

Tyrosine also increased perseveration in S1. We recently reported that L-dopa reduced directed (uncertainty-based) exploration in a restless bandit task (Chakroun et al., 2020). This was mirrored by reduced tracking of overall uncertainty in anterior cingulate and insula cortex. As the present model did not explicitly account for exploration, the observed increases in perseveration might be due to reduced exploration. Further modeling work is required to disentangle perseveration and exploration effects in the seq. RL task.

Modeling further revealed a reduced asymptote of the drift-rate in 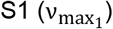, an increase in the learning rate of model-free Q-values in S1 (η_1_) and attenuated drift-rate scaling in S2 (β_2_). These findings all suggest a modulation of value updating and related drift rate modulation during RL following tyrosine. This resonates with earlier work that has linked dopaminergic transmission to value updating during RL (Diederen et al., 2017; Mathar et al., 2017; Pessiglione et al., 2006).

### Tyrosine effects on temporal discounting

In contrast to the consistent effects of tyrosine on model-based control, evidence for a reduction in temporal discounting following tyrosine supplementation was somewhat more mixed. A model agnostic mixed effects regression analysis revealed a significant interaction effect of delay*tyrosine on LL choice proportions, whereas log(*k*) in the DDM was only numerically reduced. While some studies showed similar effects following pharmacological manipulations to foster dopamine transmission (de Wit et al., 2002; Wagner et al., 2020) other studies have reported contrasting effects (Pine et al., 2010; Weber et al., 2016). These inconsistencies may partly be due to potential inverted U-shaped associations between central dopamine levels and performance (Cools & D’Esposito, 2011; Petzold et al., 2019). However, tyrosine-related changes in temporal discounting were not differentially modulated by blink rate, a potential proxy measure for central dopamine levels, arguing against this account (Groman et al., 2014; Jongkees & Colzato, 2016). But see a more recent publication from Sescousse et al. (2018).

There is also growing evidence for a modulatory role of noradrenaline transmission on measures of impulsivity. In 2006, Chamberlain et al. showed for the first time that increased noradrenaline transmission via the re-uptake inhibitor Atomoxetine improved response inhibition. This finding has been replicated in attention-deficit hyperactivity disorder (Chamberlain et al., 2007) and Parkinson’s disease patients (Kehagia et al., 2014) and was also replicated using the noradrenergic antagonist Yohimbine (Herman et al., 2019). In contrast, they found no effect on temporal discounting performance. However, they used the Monetary Choice Questionnaire (Kirby et al., 1999) that only incorporates 27 choices and thus may lack sensitivity in comparison to our temporal discounting task with 128 choices that allowed subsequent modeling of choice behavior and response time distributions via hierarchical Bayesian DDM.

### Associations between physiological arousal and temporal discounting

We ran exploratory analyses to link physiological arousal to task performance. Greater mean pupil dilation at baseline (t0) predicted increased temporal discounting across model-agnostic (larger-later choice proportions) and model-based measures (log(k), bias parameter z). This extends previous findings on pupillometry correlates of temporal discounting at the trial level (Lempert et al., 2015, 2016). For example, Lempert et al., (2016) found that greater pupil dilation during processing of choice options was associated with an increased probability to select the larger-later reward. In contrast, in the present study, greater pupil dilation *before* task performance predicted *more* temporal discounting. Trial-wise pupillometry measures, reflecting transient changes in arousal thus might show reverse effects compared to trait-like arousal levels at rest. Schmidt et al. (2013) found a negative association of arousal (heart rate) at rest and risk-taking, contrasting with our finding of a positive association of arousal at rest (pupil dilation) and impulsive choice. Thus, future studies of catecholamine dependent cognitive functioning and related supplementation interventions should incorporate state of the art modeling approaches and physiological correlates of arousal to shed light on the mixed findings so far (Unsworth et al., 2019).

### Limitations

Our study has a number of limitations that need to be acknowledged. First, we only included male participants. This was done in order to minimize inter-individual variability in estrogen levels known to modulate central DA transmission with potential implications for tyrosine effects (Di Paolo, 1994; Diekhof & Ratnayake, 2016). Second, our final sample size of n=28 is relatively small. However, when compared to earlier supplementation work it still is among the largest within-subject studies on tyrosine supplementation reported so far (Jongkees et al., 2015). Future studies would nonetheless benefit from larger sample sizes and a balanced distribution of male and female participants. Third, we used a fixed single dose of 2g tyrosine per participant. While this makes the present supplementation regime comparable to a large number of previous studies (Jongkees et al, 2015), future studies could benefit from an examination of potential dose-response effects, and/or examine the potential benefits of individualized dosages. For example, van de Rest et al. (2017) found that higher doses of tyrosine (amounting to 7-14g for a participant of 70 kg) were detrimental to WM task performance compared with lower doses. However, dose-response effects at lower amounts are unclear. Likewise, long-term effects of tyrosine supplementation have so far not been examined in detail, despite promising cognitive performance findings of habitual tyrosine intake in daily nutrition (Kühn et al., 2019). In addition, it could be argued that the tests on tyrosine-related changes in model parameters, such as directional Bayes Factors, should be considered for correction of multiple comparisons. However, in a hierarchical Bayesian framework, the risk of Type S (and likewise Type M) errors is substantially ameliorated as differences in posterior means are shrinked to a greater extent than posterior standard deviations in the light of partial pooling (Gelman et al., 2012). Furthermore, we did not observe a correlation between baseline spontaneous eye blink rate as a proxy measure of baseline DA levels (Jongkees & Colzato, 2016) and tyrosine-related changes in task performance. However, future studies should consider possible modulatory effects of baseline DA or NA transmission levels (Cools & D’Esposito, 2011; Jongkees, 2020), which will likely require much larger samples. Finally, although we assessed physiological arousal markers, we did not directly measure central DA or NA transmission nor related metabolites. Thus, direct evidence for increased catecholamine neurotransmission following the present supplementation regime (2g tyrosine) is still lacking. Nonetheless, several earlier studies in rodents and humans have shown that a single dose of tyrosine can increase catecholamine release (Glaeser et al., 1979; Growdon et al., 1982; Kishore et al., 2013).

## Conclusion

Potential cognitive enhancement effects of amino acid supplementation have gained considerable interest. Here we show that a single dose of tyrosine supplementation (2g) reduced decision thresholds across two value-based decision-making tasks, temporal discounting and model-based RL. Model-based reinforcement learning was improved following tyrosine supplementation, and temporal discounting was, if anything, reduced. These findings show that model-based approaches can reveal novel insights into computational effects of supplementation. We also for the first time comprehensively report exploratory analyses of potential physiological correlates of tyrosine supplementation. These analyses revealed an overall reduction in physiological arousal following tyrosine supplementation, complementing previous approaches that focused solely on behavioral measures. Taken together, these results suggest specific computational and physiological effects of tyrosine supplementation that future studies can build upon.

## Acknowledgements

This work was funded by Deutsche Forschungsgemeinschaft (PE1627/5-1 to J.P.)

## References

Acheson, A., & de Wit, H. (2008). Bupropion improves attention but does not affect impulsive behavior in healthy young adults. Experimental and Clinical Psychopharmacology, 16(2), 113–123. https://doi.org/10.1037/1064-1297.16.2.113

Aminihajibashi, S., Hagen, T., Foldal, M. D., Laeng, B., & Espeseth, T. (2019). Individual differences in resting-state pupil size: Evidence for association between working memory capacity and pupil size variability. International Journal of Psychophysiology, 140, 1–7. https://doi.org/10.1016/j.ijpsycho.2019.03.007

Amlung, M., Marsden, E., Holshausen, K., Morris, V., Patel, H., Vedelago, L., Naish, K. R., Reed, D. D., & McCabe, R. E. (2019). Delay Discounting as a Transdiagnostic Process in Psychiatric Disorders: A Meta-analysis. JAMA Psychiatry, 76(11), 1176–1186. https://doi.org/10.1001/jamapsychiatry.2019.2102

Aron, A. R., Herz, D. M., Brown, P., Forstmann, B. U., & Zaghloul, K. (2016). Frontosubthalamic Circuits for Control of Action and Cognition. The Journal of Neuroscience: The Official Journal of the Society for Neuroscience, 36(45), 11489–11495. https://doi.org/10.1523/JNEUROSCI.2348-16.2016

Balleine, B. W., & O’Doherty, J. P. (2010). Human and rodent homologies in action control: Corticostriatal determinants of goal-directed and habitual action. Neuropsychopharmacology: Official Publication of the American College of Neuropsychopharmacology, 35(1), 48–69. https://doi.org/10.1038/npp.2009.131

Balodis, I. M., Kober, H., Worhunsky, P. D., Stevens, M. C., Pearlson, G. D., & Potenza, M. N. (2012). Diminished frontostriatal activity during processing of monetary rewards and losses in pathological gambling. Biological Psychiatry, 71(8), 749–757. https://doi.org/10.1016/j.biopsych.2012.01.006

Barbato, G., Ficca, G., Muscettola, G., Fichele, M., Beatrice, M., & Rinaldi, F. (2000). Diurnal variation in spontaneous eye-blink rate. Psychiatry Research, 93(2), 145–151. https://doi.org/10.1016/s0165-1781(00)00108-6

Bartels, C., Wagner, M., Wolfsgruber, S., Ehrenreich, H., Schneider, A., & Alzheimer’s Disease Neuroimaging Initiative. (2018). Impact of SSRI Therapy on Risk of Conversion From Mild Cognitive Impairment to Alzheimer’s Dementia in Individuals With Previous Depression. The American Journal of Psychiatry, 175(3), 232–241. https://doi.org/10.1176/appi.ajp.2017.17040404

Bayer, H. M., & Glimcher, P. W. (2005). Midbrain dopamine neurons encode a quantitative reward prediction error signal. Neuron, 47(1), 129–141. https://doi.org/10.1016/j.neuron.2005.05.020

Beard, E., Dienes, Z., Muirhead, C., & West, R. (2016). Using Bayes factors for testing hypotheses about intervention effectiveness in addictions research. Addiction (Abingdon, England), 111(12), 2230–2247. https://doi.org/10.1111/add.13501

Beck, A. T., Steer, R. A., Ball, R., & Ranieri, W. (1996). Comparison of Beck Depression Inventories -IA and -II in psychiatric outpatients. Journal of Personality Assessment, 67(3), 588–597. https://doi.org/10.1207/s15327752jpa6703_13

Belujon, P., & Grace, A. A. (2017). Dopamine System Dysregulation in Major Depressive Disorders. The International Journal of Neuropsychopharmacology, 20(12), 1036–1046. https://doi.org/10.1093/ijnp/pyx056

Benjamini, Y., & Hochberg, Y. (1995). Controlling the False Discovery Rate: A Practical and Powerful Approach to Multiple Testing. Journal of the Royal Statistical Society. Series B (Methodological), 57(1), 289–300.

Berntson, G. G., Thomas Bigger Jr., J., Eckberg, D. L., Grossman, P., Kaufmann, P. G., Malik, M., Nagaraja, H. N., Porges, S. W., Saul, J. P., Stone, P. H., & Van Der Molen, M. W. (1997). Heart rate variability: Origins, methods, and interpretive caveats. Psychophysiology, 34(6), 623–648. https://doi.org/10.1111/j.1469-8986.1997.tb02140.x

Bickel, W. K., Jarmolowicz, D. P., Mueller, E. T., Koffarnus, M. N., & Gatchalian, K. M. (2012). Excessive discounting of delayed reinforcers as a trans-disease process contributing to addiction and other disease-related vulnerabilities: Emerging evidence. Pharmacology & Therapeutics, 134(3), 287–297. https://doi.org/10.1016/j.pharmthera.2012.02.004

Bradley, M. M., Miccoli, L., Escrig, M. A., & Lang, P. J. (2008). The pupil as a measure of emotional arousal and autonomic activation. Psychophysiology, 45(4), 602–607. https://doi.org/10.1111/j.1469-8986.2008.00654.x

Brodnik, Z., Bongiovanni, R., Double, M., & Jaskiw, G. E. (2012). Increased tyrosine availability increases brain regional DOPA levels in vivo. Neurochemistry International, 61(7), 1001–1006. https://doi.org/10.1016/j.neuint.2012.07.012

Bromberg-Martin, E. S., Matsumoto, M., & Hikosaka, O. (2010). Dopamine in motivational control: Rewarding, aversive, and alerting. Neuron, 68(5), 815–834. https://doi.org/10.1016/j.neuron.2010.11.022

Bruder, L. R., Scharer, L., & Peters, J. (2021). Reliability assessment of temporal discounting measures in virtual reality environments. Scientific Reports, 11, 7015. https://doi.org/10.1038/s41598-021-86388-8

Canteras, N. S., Shammah-Lagnado, S. J., Silva, B. A., & Ricardo, J. A. (1990). Afferent connections of the subthalamic nucleus: A combined retrograde and anterograde horseradish peroxidase study in the rat. Brain Research, 513(1), 43–59. https://doi.org/10.1016/0006-8993(90)91087-w

Carver, C. S., & White, T. L. (1994). Behavioral inhibition, behavioral activation, and affective responses to impending reward and punishment: The BIS/BAS Scales. Journal of Personality and Social Psychology, 67(2), 319–333. https://doi.org/10.1037/0022-3514.67.2.319

Chakroun, K., Mathar, D., Wiehler, A., Ganzer, F., & Peters, J. (2020). Dopaminergic modulation of the exploration/exploitation trade-off in human decision-making. ELife, 9. https://doi.org/10.7554/eLife.51260

Chamberlain, S. R., Müller, U., Blackwell, A. D., Clark, L., Robbins, T. W., & Sahakian, B. J. (2006). Neurochemical Modulation of Response Inhibition and Probabilistic Learning in Humans. Science, 311(5762), 861–863. https://doi.org/10.1126/science.1121218

Chamberlain, S. R., Robbins, T. W., & Sahakian, B. J. (2007). The neurobiology of attention-deficit/hyperactivity disorder. Biological Psychiatry, 61(12), 1317–1319. https://doi.org/10.1016/j.biopsych.2007.04.009

Chopra, A. S., Lordan, R., Horbañczuk, O. K., Atanasov, A. G., Chopra, I., Horbañczuk, J. O., Jóźwik, A., Huang, L., Pirgozliev, V., Banach, M., Battino, M., & Arkells, N. (2022). The current use and evolving landscape of nutraceuticals. Pharmacological Research, 175, 106001. https://doi.org/10.1016/j.phrs.2021.106001

Colzato, L. S., Jongkees, B. J., Sellaro, R., & Hommel, B. (2013). Working memory reloaded: Tyrosine repletes updating in the N-back task. Frontiers in Behavioral Neuroscience, 7, 200. https://doi.org/10.3389/fnbeh.2013.00200

Colzato, L. S., Jongkees, B. J., Sellaro, R., van den Wildenberg, W. P. M., & Hommel, B. (2014). Eating to stop: Tyrosine supplementation enhances inhibitory control but not response execution. Neuropsychologia, 62, 398–402. https://doi.org/10.1016/j.neuropsychologia.2013.12.027

Cools, R., & D’Esposito, M. (2011). Inverted-U-shaped dopamine actions on human working memory and cognitive control. Biological Psychiatry, 69(12), e113–125. https://doi.org/10.1016/j.biopsych.2011.03.028

Cragg, S. J., Baufreton, J., Xue, Y., Bolam, J. P., & Bevan, M. D. (2004). Synaptic release of dopamine in the subthalamic nucleus. The European Journal of Neuroscience, 20(7), 1788–1802. https://doi.org/10.1111/j.1460-9568.2004.03629.x

Culbreth, A. J., Westbrook, A., Daw, N. D., Botvinick, M., & Barch, D. M. (2016). Reduced model-based decision-making in schizophrenia. Journal of Abnormal Psychology, 125(6), 777–787. https://doi.org/10.1037/abn0000164

Daw, N. D., Gershman, S. J., Seymour, B., Dayan, P., & Dolan, R. J. (2011). Model-based influences on humans’ choices and striatal prediction errors. Neuron, 69(6), 1204–1215. https://doi.org/10.1016/j.neuron.2011.02.027

de Wit, H., Enggasser, J. L., & Richards, J. B. (2002). Acute administration of d-amphetamine decreases impulsivity in healthy volunteers. Neuropsychopharmacology: Official Publication of the American College of Neuropsychopharmacology, 27(5), 813–825. https://doi.org/10.1016/S0893-133X(02)00343-3

Deijen, J. B., Wientjes, C. J. E., Vullinghs, H. F. M., Cloin, P. A., & Langefeld, J. J. (1999). Tyrosine improves cognitive performance and reduces blood pressure in cadets after one week of a combat training course. Brain Research Bulletin, 48(2), 203–209. https://doi.org/10.1016/S0361-9230(98)00163-4

Deserno, L. (2015). Lateral prefrontal model-based signatures are reduced in healthy individuals with high trait impulsivity. Translational Psychiatry, 9.

Di Paolo, T. (1994). Modulation of brain dopamine transmission by sex steroids. Reviews in the Neurosciences, 5(1), 27–41. https://doi.org/10.1515/revneuro.1994.5.1.27

Diederen, K. M. J., Ziauddeen, H., Vestergaard, M. D., Spencer, T., Schultz, W., & Fletcher, P. C. (2017). Dopamine Modulates Adaptive Prediction Error Coding in the Human Midbrain and Striatum. The Journal of Neuroscience: The Official Journal of the Society for Neuroscience, 37(7), 1708–1720. https://doi.org/10.1523/JNEUROSCI.1979-16.2016

Diekhof, E. K., & Ratnayake, M. (2016). Menstrual cycle phase modulates reward sensitivity and performance monitoring in young women: Preliminary fMRI evidence. Neuropsychologia, 84, 70–80. https://doi.org/10.1016/j.neuropsychologia.2015.10.016

Dolan, R. J., & Dayan, P. (2013). Goals and habits in the brain. Neuron, 80(2), 312–325. https://doi.org/10.1016/j.neuron.2013.09.007

Doll, B. B., Simon, D. A., & Daw, N. D. (2012). The ubiquity of model-based reinforcement learning. Current Opinion in Neurobiology, 22(6), 1075–1081. https://doi.org/10.1016/j.conb.2012.08.003

Elsworth, J. D., Lawrence, M. S., Roth, R. H., Taylor, J. R., Mailman, R. B., Nichols, D. E., Lewis, M. H., & Redmond, D. E. (1991). D1 and D2 dopamine receptors independently regulate spontaneous blink rate in the vervet monkey. The Journal of Pharmacology and Experimental Therapeutics, 259(2), 595–600.

Enkavi, A. Z., Eisenberg, I. W., Bissett, P. G., Mazza, G. L., MacKinnon, D. P., Marsch, L. A., & Poldrack, R. A. (2019). Large-scale analysis of test-retest reliabilities of self-regulation measures. Proceedings of the National Academy of Sciences of the United States of America, 116(12), 5472–5477. https://doi.org/10.1073/pnas.1818430116

Foerde, K., Figner, B., Doll, B. B., Woyke, I. C., Braun, E. K., Weber, E. U., & Shohamy, D. (2016). Dopamine Modulation of Intertemporal Decision-making: Evidence from Parkinson Disease. Journal of Cognitive Neuroscience, 28(5), 657–667. https://doi.org/10.1162/jocn_a_00929

Fontanesi, L., Gluth, S., Spektor, M. S., & Rieskamp, J. (2019). A reinforcement learning diffusion decision model for value-based decisions. Psychonomic Bulletin & Review, 26(4), 1099–1121. https://doi.org/10.3758/s13423-018-1554-2

Frank, M. J., Samanta, J., Moustafa, A. A., & Sherman, S. J. (2007). Hold your horses: Impulsivity, deep brain stimulation, and medication in parkinsonism. Science (New York, N.Y.), 318(5854), 1309–1312. https://doi.org/10.1126/science.1146157

Gelman, A., Hill, J., & Yajima, M. (2012). Why We (Usually) Don’t Have to Worry About Multiple Comparisons. Journal of Research on Educational Effectiveness, 5(2), 189–211. https://doi.org/10.1080/19345747.2011.618213

Gillan, C. M., Kosinski, M., Whelan, R., Phelps, E. A., & Daw, N. D. (o. J.). Characterizing a psychiatric symptom dimension related to deficits in goal-directed control. eLife, 5. https://doi.org/10.7554/eLife.11305

Glaeser, B. S., Melamed, E., Growdon, J. H., & Wurtman, R. J. (1979). Elevation of plasma tyrosine after a single oral dose of L-tyrosine. Life Sciences, 25(3), 265–271. https://doi.org/10.1016/0024-3205(79)90294-7

Gordan, R., Gwathmey, J. K., & Xie, L.-H. (2015). Autonomic and endocrine control of cardiovascular function. World Journal of Cardiology, 7(4), 204–214. https://doi.org/10.4330/wjc.v7.i4.204

Green, L., Myerson, J., & McFadden, E. (1997). Rate of temporal discounting decreases with amount of reward. Memory & Cognition, 25(5), 715–723. https://doi.org/10.3758/bf03211314

Groman, S. M., James, A. S., Seu, E., Tran, S., Clark, T. A., Harpster, S. N., Crawford, M., Burtner, J. L., Feiler, K., Roth, R. H., Elsworth, J. D., London, E. D., & Jentsch, J. D. (2014). In the blink of an eye: Relating positive-feedback sensitivity to striatal dopamine D2-like receptors through blink rate. The Journal of Neuroscience: The Official Journal of the Society for Neuroscience, 34(43), 14443–14454. https://doi.org/10.1523/JNEUROSCI.3037-14.2014

Growdon, J. H., Melamed, E., Logue, M., Hefti, F., & Wurtman, R. J. (1982). Effects of oral L-tyrosine administration on CSF tyrosine and homovanillic acid levels in patients with Parkinson’s disease. Life Sciences, 30(10), 827–832. https://doi.org/10.1016/0024-3205(82)90596-3

Guitart-Masip, M., Beierholm, U. R., Dolan, R., Duzel, E., & Dayan, P. (2011). Vigor in the face of fluctuating rates of reward: An experimental examination. Journal of Cognitive Neuroscience, 23(12), 3933–3938. https://doi.org/10.1162/jocn_a_00090

Hase, A., Jung, S. E., & aan het Rot, M. (2015). Behavioral and cognitive effects of tyrosine intake in healthy human adults. Pharmacology Biochemistry and Behavior, 133, 1–6. https://doi.org/10.1016/j.pbb.2015.03.008

Herman, A. M., Critchley, H. D., & Duka, T. (2019). The impact of Yohimbine-induced arousal on facets of behavioural impulsivity. Psychopharmacology, 236(6), 1783–1795. https://doi.org/10.1007/s00213-018-5160-9

Herz, D. M., Zavala, B. A., Bogacz, R., & Brown, P. (2016). Neural Correlates of Decision Thresholds in the Human Subthalamic Nucleus. Current Biology: CB, 26(7), 916–920. https://doi.org/10.1016/j.cub.2016.01.051

Hofmans, L., Westbrook, A., van den Bosch, R., Booij, J., Verkes, R.-J., & Cools, R. (2021). Effects of average reward rate on vigor as a function of individual variation in striatal dopamine. Psychopharmacology. https://doi.org/10.1007/s00213-021-06017-0

Holland, N., Robbins, T. W., & Rowe, J. B. (2021). The role of noradrenaline in cognition and cognitive disorders. Brain, awab111. https://doi.org/10.1093/brain/awab111

Jongkees, B. J. (2020). Baseline-dependent effect of dopamine’s precursor L-tyrosine on working memory gating but not updating. Cognitive, Affective & Behavioral Neuroscience. https://doi.org/10.3758/s13415-020-00783-8

Jongkees, B. J., & Colzato, L. S. (2016). Spontaneous eye blink rate as predictor of dopamine-related cognitive function-A review. Neuroscience and Biobehavioral Reviews, 71, 58–82. https://doi.org/10.1016/j.neubiorev.2016.08.020

Jongkees, B. J., Hommel, B., Kühn, S., & Colzato, L. S. (2015). Effect of tyrosine supplementation on clinical and healthy populations under stress or cognitive demands—A review. Journal of Psychiatric Research, 70, 50–57. https://doi.org/10.1016/j.jpsychires.2015.08.014

Joutsa, J., Voon, V., Johansson, J., Niemelä, S., Bergman, J., & Kaasinen, V. (2015). Dopaminergic function and intertemporal choice. Translational Psychiatry, 5, e491. https://doi.org/10.1038/tp.2014.133

Kaminer, J., Powers, A. S., Horn, K. G., Hui, C., & Evinger, C. (2011). Characterizing the Spontaneous Blink Generator: An Animal Model. The Journal of Neuroscience, 31(31), 11256–11267. https://doi.org/10.1523/JNEUROSCI.6218-10.2011

Kayser, A. (2019). Dopamine and Gambling Disorder: Prospects for Personalized Treatment. Current Addiction Reports, 6(2), 65–74. https://doi.org/10.1007/s40429-019-00240-8

Kehagia, A. A., Housden, C. R., Regenthal, R., Barker, R. A., Müller, U., Rowe, J., Sahakian, B. J., & Robbins, T. W. (2014). Targeting impulsivity in Parkinson’s disease using atomoxetine. Brain: A Journal of Neurology, 137(Pt 7), 1986–1997. https://doi.org/10.1093/brain/awu117

Kirby, K. N. (2009). One-year temporal stability of delay-discount rates. Psychonomic Bulletin & Review, 16(3), 457–462. https://doi.org/10.3758/PBR.16.3.457

Kishore, K., Ray, K., Anand, J. P., Thakur, L., Kumar, S., & Panjwani, U. (2013). Tyrosine ameliorates heat induced delay in event related potential P300 and contingent negative variation. Brain and Cognition, 83(3), 324–329. https://doi.org/10.1016/j.bandc.2013.09.005

Kool, W., Cushman, F. A., & Gershman, S. J. (2016). When Does Model-Based Control Pay Off? PLoS Computational Biology, 12(8), e1005090. https://doi.org/10.1371/journal.pcbi.1005090

Kroemer, N. B., Lee, Y., Pooseh, S., Eppinger, B., Goschke, T., & Smolka, M. N. (2019). L-DOPA reduces model-free control of behavior by attenuating the transfer of value to action. NeuroImage, 186, 113–125. https://doi.org/10.1016/j.neuroimage.2018.10.075

Kühn, S., Düzel, S., Colzato, L., Norman, K., Gallinat, J., Brandmaier, A. M., Lindenberger, U., & Widaman, K. F. (2019). Food for thought: Association between dietary tyrosine and cognitive performance in younger and older adults. Psychological Research, 83(6), 1097–1106. https://doi.org/10.1007/s00426-017-0957-4

Kühner, C., Bürger, C., Keller, F., & Hautzinger, M. (2007). Reliabilität und Validität des revidierten Beck-Depressionsinventars (BDI-II). Der Nervenarzt, 78(6), 651–656. https://doi.org/10.1007/s00115-006-2098-7

Le Masurier, M., Oldenzeil, W., Lehman, C., Cowen, P., & Sharp, T. (2006). Effect of acute tyrosine depletion in using a branched chain amino-acid mixture on dopamine neurotransmission in the rat brain. Neuropsychopharmacology: Official Publication of the American College of Neuropsychopharmacology, 31(2), 310–317. https://doi.org/10.1038/sj.npp.1300835

Lempert, K. M., Glimcher, P. W., & Phelps, E. A. (2015). Emotional arousal and discount rate in intertemporal choice are reference dependent. Journal of Experimental Psychology. General, 144(2), 366–373. https://doi.org/10.1037/xge0000047

Lempert, K. M., Johnson, E., & Phelps, E. A. (2016). Emotional arousal predicts intertemporal choice. Emotion (Washington, D.C.), 16(5), 647–656. https://doi.org/10.1037/emo0000168

Lempert, K. M., Steinglass, J. E., Pinto, A., Kable, J. W., & Simpson, H. B. (2019). Can delay discounting deliver on the promise of RDoC? Psychological Medicine, 49(2), 190–199. https://doi.org/10.1017/S0033291718001770

Magill, R. A., Waters, W. F., Bray, G. A., Volaufova, J., Smith, S. R., Lieberman, H. R., McNevin, N., & Ryan, D. H. (2003). Effects of tyrosine, phentermine, caffeine D-amphetamine, and placebo on cognitive and motor performance deficits during sleep deprivation. Nutritional Neuroscience, 6(4), 237–246. https://doi.org/10.1080/1028415031000120552

Mathar, D., Wilkinson, L., Holl, A. K., Neumann, J., Deserno, L., Villringer, A., Jahanshahi, M., & Horstmann, A. (2017). The role of dopamine in positive and negative prediction error utilization during incidental learning—Insights from Positron Emission Tomography, Parkinson’s disease and Huntington’s disease. Cortex; a Journal Devoted to the Study of the Nervous System and Behavior, 90, 149–162. https://doi.org/10.1016/j.cortex.2016.09.004

Meule, A., Michalek, S., Friederich, H.-C., & Brockmeyer, T. (2020). Confirmatory factor analysis of the Barratt Impulsiveness Scale-short form (BIS-15) in patients with mental disorders. Psychiatry Research, 284, 112665. https://doi.org/10.1016/j.psychres.2019.112665

Moja, E. A., Lucini, V., Benedetti, F., & Lucca, A. (1996). Decrease in plasma phenylalanine and tyrosine after phenylalanine-tyrosine free amino acid solutions in man. Life Sciences, 58(26), 2389–2395. https://doi.org/10.1016/0024-3205(96)00242-1

Molinoff, P. B., & Axelrod, J. (1971). Biochemistry of catecholamines. Annual Review of Biochemistry, 40, 465–500. https://doi.org/10.1146/annurev.bi.40.070171.002341

Murphy, P. R., O’Connell, R. G., O’Sullivan, M., Robertson, I. H., & Balsters, J. H. (2014). Pupil diameter covaries with BOLD activity in human locus coeruleus. Human Brain Mapping, 35(8), 4140–4154. https://doi.org/10.1002/hbm.22466

Nambu, A., Tokuno, H., & Takada, M. (2002). Functional significance of the cortico-subthalamo-pallidal „hyperdirect” pathway. Neuroscience Research, 43(2), 111–117. https://doi.org/10.1016/s0168-0102(02)00027-5

O’Brien, C., Mahoney, C., Tharion, W. J., Sils, I. V., & Castellani, J. W. (2007). Dietary tyrosine benefits cognitive and psychomotor performance during body cooling. Physiology & Behavior, 90(2), 301–307. https://doi.org/10.1016/j.physbeh.2006.09.027

Parent, A., & Hazrati, L. N. (1995). Functional anatomy of the basal ganglia. II. The place of subthalamic nucleus and external pallidum in basal ganglia circuitry. Brain Research. Brain Research Reviews, 20(1), 128–154. https://doi.org/10.1016/0165-0173(94)00008-d

Pedersen, M. L., Frank, M. J., & Biele, G. (2017). The drift diffusion model as the choice rule in reinforcement learning. Psychonomic Bulletin & Review, 24(4), 1234–1251. https://doi.org/10.3758/s13423-016-1199-y

Pessiglione, M., Seymour, B., Flandin, G., Dolan, R. J., & Frith, C. D. (2006). Dopamine-dependent prediction errors underpin reward-seeking behaviour in humans. Nature, 442(7106), 1042–1045. https://doi.org/10.1038/nature05051

Peters, J., & Büchel, C. (2011). The neural mechanisms of inter-temporal decision-making: Understanding variability. Trends in Cognitive Sciences, 15(5), 227–239. https://doi.org/10.1016/j.tics.2011.03.002

Peters, J., & D’Esposito, M. (2020a). The drift diffusion model as the choice rule in inter-temporal and risky choice: A case study in medial orbitofrontal cortex lesion patients and controls. PLOS Computational Biology, 16(4), e1007615. https://doi.org/10.1371/journal.pcbi.1007615

Peters, J., & D’Esposito, M. (2020b). The drift diffusion model as the choice rule in inter-temporal and risky choice: A case study in medial orbitofrontal cortex lesion patients and controls. PLoS Computational Biology, 16(4). https://doi.org/10.1371/journal.pcbi.1007615

Petzold, J., Kienast, A., Lee, Y., Pooseh, S., London, E. D., Goschke, T., & Smolka, M. N. (2019). Baseline impulsivity may moderate L-DOPA effects on value-based decision-making. Scientific Reports, 9(1), 5652. https://doi.org/10.1038/s41598-019-42124-x

Phillips, M. A., Szabadi, E., & Bradshaw, C. M. (2000). Comparison of the effects of clonidine and yohimbine on spontaneous pupillary fluctuations in healthy human volunteers. Psychopharmacology, 150(1), 85–89. https://doi.org/10.1007/s002130000398

Pine, A., Shiner, T., Seymour, B., & Dolan, R. J. (2010). Dopamine, time, and impulsivity in humans. The Journal of Neuroscience: The Official Journal of the Society for Neuroscience, 30(26), 8888–8896. https://doi.org/10.1523/JNEUROSCI.6028-09.2010

Preuschoff, K., ‘t Hart, B. M., & Einhäuser, W. (2011). Pupil Dilation Signals Surprise: Evidence for Noradrenaline’s Role in Decision Making. Frontiers in Neuroscience, 5, 115. https://doi.org/10.3389/fnins.2011.00115

Ratcliff, R., & McKoon, G. (2008). The diffusion decision model: Theory and data for two-choice decision tasks. Neural Computation, 20(4), 873–922. https://doi.org/10.1162/neco.2008.12-06-420

Rodríguez-Liñares, L., Méndez, A. J., Lado, M. J., Olivieri, D. N., Vila, X. A., & Gómez-Conde, I. (2011). An open source tool for heart rate variability spectral analysis. Computer Methods and Programs in Biomedicine, 103(1), 39–50. https://doi.org/10.1016/j.cmpb.2010.05.012

Roiser, J. P., McLean, A., Ogilvie, A. D., Blackwell, A. D., Bamber, D. J., Goodyer, I., Jones, P. B., & Sahakian, B. J. (2005). The subjective and cognitive effects of acute phenylalanine and tyrosine depletion in patients recovered from depression. Neuropsychopharmacology: Official Publication of the American College of Neuropsychopharmacology, 30(4), 775–785. https://doi.org/10.1038/sj.npp.1300659

Schmidt, B., Mussel, P., & Hewig, J. (2013). I’m too calm—Let’s take a risk! On the impact of state and trait arousal on risk taking. Psychophysiology, 50(5), 498–503. https://doi.org/10.1111/psyp.12032

Schneider, M., Hathway, P., Leuchs, L., Sämann, P. G., Czisch, M., & Spoormaker, V. I. (2016). Spontaneous pupil dilations during the resting state are associated with activation of the salience network. NeuroImage, 139, 189–201. https://doi.org/10.1016/j.neuroimage.2016.06.011

Schultz, W. (2011). Potential vulnerabilities of neuronal reward, risk, and decision mechanisms to addictive drugs. Neuron, 69(4), 603–617. https://doi.org/10.1016/j.neuron.2011.02.014

Schultz, W., Dayan, P., & Montague, P. R. (1997). A neural substrate of prediction and reward. Science (New York, N.Y.), 275(5306), 1593–1599. https://doi.org/10.1126/science.275.5306.1593

Sescousse, G., Ligneul, R., van Holst, R. J., Janssen, L. K., de Boer, F., Janssen, M., Berry, A. S., Jagust, W. J., & Cools, R. (2018). Spontaneous eye blink rate and dopamine synthesis capacity: Preliminary evidence for an absence of positive correlation. The European Journal of Neuroscience, 47(9), 1081–1086. https://doi.org/10.1111/ejn.13895

Shahar, N., Hauser, T. U., Moutoussis, M., Moran, R., Keramati, M., NSPN consortium, & Dolan, R. J. (2019). Improving the reliability of model-based decision-making estimates in the two-stage decision task with reaction-times and drift-diffusion modeling. PLOS Computational Biology, 15(2), e1006803. https://doi.org/10.1371/journal.pcbi.1006803

Sharp, M. E., Foerde, K., Daw, N. D., & Shohamy, D. (2016). Dopamine selectively remediates ‘model-based’ reward learning: A computational approach. Brain, 139(2), 355–364. https://doi.org/10.1093/brain/awv347

Smith, P. L., & Ratcliff, R. (2004). Psychology and neurobiology of simple decisions. Trends in Neurosciences, 27(3), 161–168. https://doi.org/10.1016/j.tins.2004.01.006

Spinella, M. (2007). Normative data and a short form of the Barratt Impulsiveness Scale. The International Journal of Neuroscience, 117(3), 359–368. https://doi.org/10.1080/00207450600588881

Steenbergen, L., Sellaro, R., Hommel, B., & Colzato, L. S. (2015). Tyrosine promotes cognitive flexibility: Evidence from proactive vs. reactive control during task switching performance. Neuropsychologia, 69, 50–55. https://doi.org/10.1016/j.neuropsychologia.2015.01.022

Steinberg, E. E., Keiflin, R., Boivin, J. R., Witten, I. B., Deisseroth, K., & Janak, P. H. (2013). A Causal Link Between Prediction Errors, Dopamine Neurons and Learning. Nature neuroscience, 16(7), 966–973. https://doi.org/10.1038/nn.3413

Strobel, A., Beauducel, A., Debener, S., & Brocke, B. (2001). Eine deutschsprachige Version des BIS/BAS-Fragebogens von Carver und White. [A German version of Carver and White’s BIS/BAS scales.]. Zeitschrift für Differentielle und Diagnostische Psychologie, 22(3), 216–227. https://doi.org/10.1024/0170-1789.22.3.216

Sved, A. F., Fernstrom, J. D., & Wurtman, R. J. (1979). Tyrosine administration reduces blood pressure and enhances brain norepinephrine release in spontaneously hypertensive rats. Proceedings of the National Academy of Sciences of the United States of America, 76(7), 3511–3514.

Toyama, A., Katahira, K., & Ohira, H. (2017). A simple computational algorithm of model-based choice preference. Cognitive, Affective & Behavioral Neuroscience, 17(4), 764–783. https://doi.org/10.3758/s13415-017-0511-2

Toyama, A., Katahira, K., & Ohira, H. (2019). Reinforcement Learning With Parsimonious Computation and a Forgetting Process. Frontiers in Human Neuroscience, 13, 153. https://doi.org/10.3389/fnhum.2019.00153

Unsworth, N., Robison, M. K., & Miller, A. L. (2019). Individual differences in baseline oculometrics: Examining variation in baseline pupil diameter, spontaneous eye blink rate, and fixation stability. Cognitive, Affective, & Behavioral Neuroscience, 19(4), 1074–1093. https://doi.org/10.3758/s13415-019-00709-z

van de Rest, O., Bloemendaal, M., de Heus, R., & Aarts, E. (2017). Dose-Dependent Effects of Oral Tyrosine Administration on Plasma Tyrosine Levels and Cognition in Aging. Nutrients, 9(12), 1279. https://doi.org/10.3390/nu9121279

Vehtari, A., Gelman, A., & Gabry, J. (2017). Practical Bayesian model evaluation using leave-one-out cross-validation and WAIC. Statistics and Computing, 27(5), 1413–1432. https://doi.org/10.1007/s11222-016-9696-4

Vikbladh, O. M., Meager, M. R., King, J., Blackmon, K., Devinsky, O., Shohamy, D., Burgess, N., & Daw, N. D. (2019). Hippocampal Contributions to Model-Based Planning and Spatial Memory. Neuron, 102(3), 683-693.e4. https://doi.org/10.1016/j.neuron.2019.02.014

Wagner, B., Clos, M., Sommer, T., & Peters, J. (2020). Dopaminergic Modulation of Human Intertemporal Choice: A Diffusion Model Analysis Using the D2-Receptor Antagonist Haloperidol. The Journal of Neuroscience: The Official Journal of the Society for Neuroscience, 40(41), 7936–7948. https://doi.org/10.1523/JNEUROSCI.0592-20.2020

Wagner, B., Mathar, D., & Peters, J. (2021). Gambling environment exposure increases temporal discounting but improves model-based control in regular slot-machine gamblers (S. 2021.07.15.452520). https://doi.org/10.1101/2021.07.15.452520

Weber, S. C., Beck-Schimmer, B., Kajdi, M.-E., Müller, D., Tobler, P. N., & Quednow, B. B. (2016). Dopamine D2/3- and μ-opioid receptor antagonists reduce cue-induced responding and reward impulsivity in humans. Translational Psychiatry, 6(7), e850. https://doi.org/10.1038/tp.2016.113

Wessel, J. R., Ghahremani, A., Udupa, K., Saha, U., Kalia, S. K., Hodaie, M., Lozano, A. M., Aron, A. R., & Chen, R. (2016). Stop-related subthalamic beta activity indexes global motor suppression in Parkinson’s disease. Movement Disorders: Official Journal of the Movement Disorder Society, 31(12), 1846–1853. https://doi.org/10.1002/mds.26732

Wunderlich, K., Smittenaar, P., & Dolan, R. J. (2012). Dopamine enhances model-based over model-free choice behavior. Neuron, 75(3), 418–424. https://doi.org/10.1016/j.neuron.2012.03.042

Wyckmans, F., Otto, A. R., Sebold, M., Daw, N., Bechara, A., Saeremans, M., Kornreich, C., Chatard, A., Jaafari, N., & Noël, X. (2019). Reduced model-based decision-making in gambling disorder. Scientific Reports, 9(1), 19625. https://doi.org/10.1038/s41598-019-56161-z

Yamori, Y., Fujiwara, M., Horie, R., & Lovenberg, W. (1980). The hypotensive effect of centrally administered tyrosine. European Journal of Pharmacology, 68(2), 201–204. https://doi.org/10.1016/0014-2999(80)90323-4

Zaman, M. L., & Doughty, M. J. (1997). Some methodological issues in the assessment of the spontaneous eyeblink frequency in man. Ophthalmic & Physiological Optics: The Journal of the British College of Ophthalmic Opticians (Optometrists), 17(5), 421–432.

