## Supplementary Information for "A single dose of the catecholamine precursor Tyrosine reduces physiological arousal and decreases decision thresholds in reinforcement learning and temporal discounting"

### Methods

*Computational modeling and data analysis*

*Sequential RL task*

*Softmax implementation*

We first implemented a standard softmax action selection scheme to link learned Q-values with participants' choices. Softmax action selection models choice probabilities as a sigmoid function of value differences (Sutton and Barto, 1998). In this regard, S1 choice probabilities are modelled via weighting of S1 model-free and model-based *Q*-values through a softmax function. Similarly, S2 stage action selection is modelled as a function of weighted model-free *Q*-values (Eq. 1 and Eq. 2). An additional parameter $\rho$ was included to model 1st-stage choice perseveration, $rep(a)$ that is set to 1 if the previous S1 choice was the same and is zero otherwise.

$$p\left(a_{j,t} = a \middle| s_{1,t}\right) = \frac{\exp(\beta_{mb_S} * Q_{mb}(a)\ + \beta_{mf_S} * Q_{mf_{S1}}(a) + \rho_S * rep(a))}{\sum_{a'} exp(\beta_{mb_S} * Q_{mb}(a')\ + \beta_{mf_S} * Q_{mf_{S1}}(a') + \rho_S * rep(a'))} \quad (1)$$

$$p\left(a_{j,t} = a \middle| s_{2,t}\right) = \frac{\exp(\beta_{2_S} * (Q_{mf_{S2}}(a)}{\sum_{a'} \exp(\beta_{2_S} * (Q_{mf_{S2}}(a')\ )} \quad (2)$$

with:

$\beta_{mb} = \beta_{mb} + s_{\beta_{mb}} * I_t$
$\beta_{mf_S} = \beta_{mf} + s_{\beta_{mf}} * I_t$
$\rho_S = \rho + s_\rho * I_t$
$\beta_{2_S} = \beta_2 + s_{\beta_2} * I_t$

To account for potential modulatory effects of tyrosine vs. placebo supplementation, we included additive 'shift' parameters $s_x$ for each parameter $x$ that were multiplied by dummy-coded supplementation predictors $I_t\ (= 1, TYR;\ = 0, PLC)$.

*Temporal discounting task*

*Softmax implementation*

In a first modeling scheme, similar to the modeling of the seq. RL data, we used a softmax function to link subjective values of LL and SS rewards in each trial with participants' choices:

$$P(LL)_t = \frac{\exp\left((\beta + s_\beta * I_t) * SV(LL_t)\right)}{\exp\left((\beta + s_\beta * I_t) * SV(SS_t)\right) + \exp\left((\beta + s_\beta * I_t) * SV(LL_t)\right)} \quad (3)$$

Here, *SV* is the subjective value of the larger but later reward according to Eq. 1 and $\beta$ is the inverse temperature parameter, modeling choice stochasticity. $SV(SS_t)$ was fixed at 20 and $I_t$ is again the dummy-coded predictor of supplementation condition, and $s_\beta$ models a potential modulatory effect of tyrosine on $\beta$.

### Results

*Demographic & psychological screening*

| | mean (range) | se |
|---|---|---|
| age | 25.25 (20-34) | 0.75 |
| BMI | 24.13 (19.6-28.89) | 0.48 |

| | | |
|---|---|---|
| YOE | 12.5 (11-15) | 0.15 |
| income | 1023.93 (0-5000) | 169.7 |
| BDI-II | 5.11 (0-18) | 0.8 |
| BIS-15 | 34.18 (26-44) | 1.05 |
| BIS(BIS/BAS) | 2.35 (1.57-3.29) | 0.08 |
| BAS(BIS/BAS | 3.05 (2.46-3.85) | 0.06 |

Table S1. Study sample characteristics (N=28). SE= standard error; YOE=years of education; BDI-II: Beck Depression Inventory-II; BIS-15: Barratt-Impulsiveness Scale (15 items); BIS/BAS: Behavioral Inhibition and Behavioral Activation System.

| S1 - RTs | $\beta$ | SE | t | p |
|---|---|---|---|---|
| interc | .6 | 0.02 | 34.12 | **<2*10^-16** |
| rew | 8.55*10^-4 | 2.26*10^-3 | .38 | .71 |
| **trans** | **5.14*10^-3** | **1.29*10^-3** | **2.73** | **7.41*10^-3** |
| **TYR** | **-8.9*10^-3** | **1.86*10^-3** | **-4.8** | **1.61*10^-6** |
| rew*trans | 1.88*10^-3 | 1.88*10^-3 | -1.0 | .32 |
| rew*TYR | -4.43*10^-4 | 1.85*10^-3 | -.24 | .81 |
| trans*TYR | -1-53*10^-3 | 1.84*10^-3 | -.83 | .41 |
| rew*trans*TYR | -2.77*10^-3 | 1.85*10^-3 | -1.5 | .13 |

**Table S2.** Effects on participants S1 RTs from a mixed effects regression analysis (rew=reward; trans=state transition; TYR=tyrosine).

*Model-comparison, predictive accuracy*

Depicted below is the model fit comparison according to Watanabe-Akaike Information Criterion (WAIC) and the estimated log pointwise predictive density (elpd).

| | drift-rate modulation | WAIC | -elpd | -Δelpd | 95% CI (-Δelpd) |
|---|---|---|---|---|---|
| Temp. Discount. | | | | | |
| **$DDM_0$** | - | 10946 | 5518 | 2027 | 1755 - 2298 |
| **$DDM_{lin}$** | linear | 8176 | 4141 | 650 | 513 - 788 |
| **$DDM_s$** | sigmoid | 6880 | 3491 | - | - |
| *Seq. RL* | | | | | |
| **$DDM_0$** | - | 20989 | 10548 | 8932 | 7997 - 9868 |
| **$DDM_{lin}$** | linear | 3943 | 2021 | 405 | 244 - 567 |
| **$DDM_s$** | sigmoid | 3136 | 1616 | - | - |

**Table S3.** Model fit comparison of the DDMs in the temporal discounting & the seq. RL task via the Watanabe-Akaike Information Criterion (WAIC), the estimated log pointwise predictive density (elpd), and its difference to the winning model ($DDM_s$).

We compared the respective model formulations also in terms of predictive accuracy of participants' binary choices in each task. While the SM model formulation performed best, as it is only fitted to participants' choices (and not to RT distributions), the $DDM_s$ performed on a comparable level and substantially better than the $DDM_0$ and $DDM_{lin}$ (Table S1).

| | softmax | $DDM_0$ | $DDM_{lin}$ | $DDM_s$ |
|---|---|---|---|---|
| **TD task** | 87 (66-93) | 64 (49-100) | 76 (60-87) | 83 (64-96) |
| **seq. RL, S1** | 80 (50-96) | 56 (49-77) | 73 (51-92) | 75 (51-95) |
| **seq. RL, S2** | 81 (55-94) | 51 (47-57) | 72 (51-87) | 75 (52-90) |

**Table S4**. Proportions of correctly predicted binary choices (mean (range)) for the temporal discounting (TD) task data and for both stages (S1, S2) of the seq. RL choice data, respectively. Values are computed via simulations based on 500 samples drawn from each of the respective single subject parameters' posterior distributions.

*Posterior predictive checks*

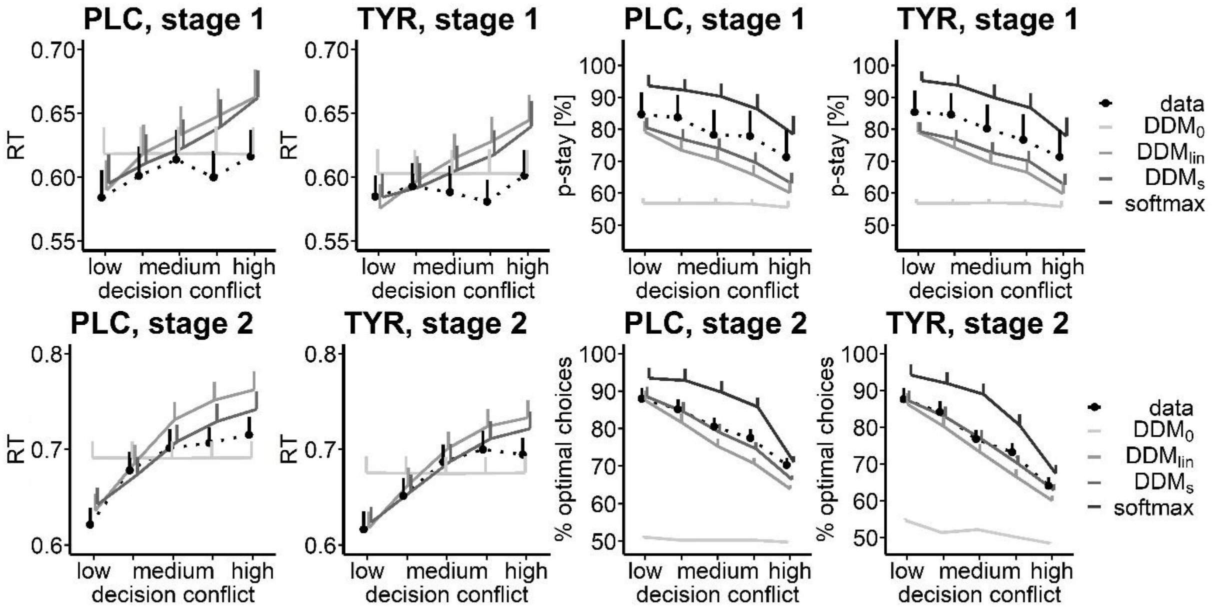


**Figure S1.** Posterior predictive checks for the winning DDMs and alternative model formulations for the seq. RL task data for the placebo and tyrosine condition. In each plot the dotted line depicts participants' median RTs or mean choice behavior. Solid lines depict the median RTs and mean choice behavior drawn from 500 simulations of each of the different DDM formulations, as well as of a standard softmax model for choice data. The upper row depicts RT data and simulations and participants' probability to choose the same action as in the previous trial for the first decision stage S1 in relation to the options value differences ('decision conflict'). The lower row depicts S2 RT data and fraction of optimal choices in S2 (highest value option chosen) of participants and related model simulations.

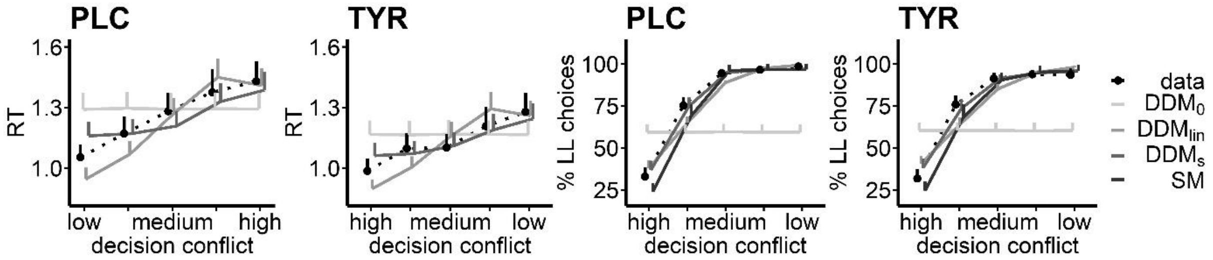


**Figure S2.** Posterior predictive checks for the winning DDMs and alternative model formulations for the temporal discounting task data for the placebo and tyrosine condition. In each plot the dotted line depicts participants' median RTs or mean % LL choices. Solid lines depict the median RTs or mean LL choices drawn from 500 simulations of each of the different DDM formulations, as well as of a standard softmax model for choice data. Data and simulations are plotted in relation to the absolute difference of LL (subjective values) and SS choice options.

*Sequential RL task, drift diffusion model*

Depicted below are the posterior distributions of the group-level mean parameter from the DDMs for the placebo condition.

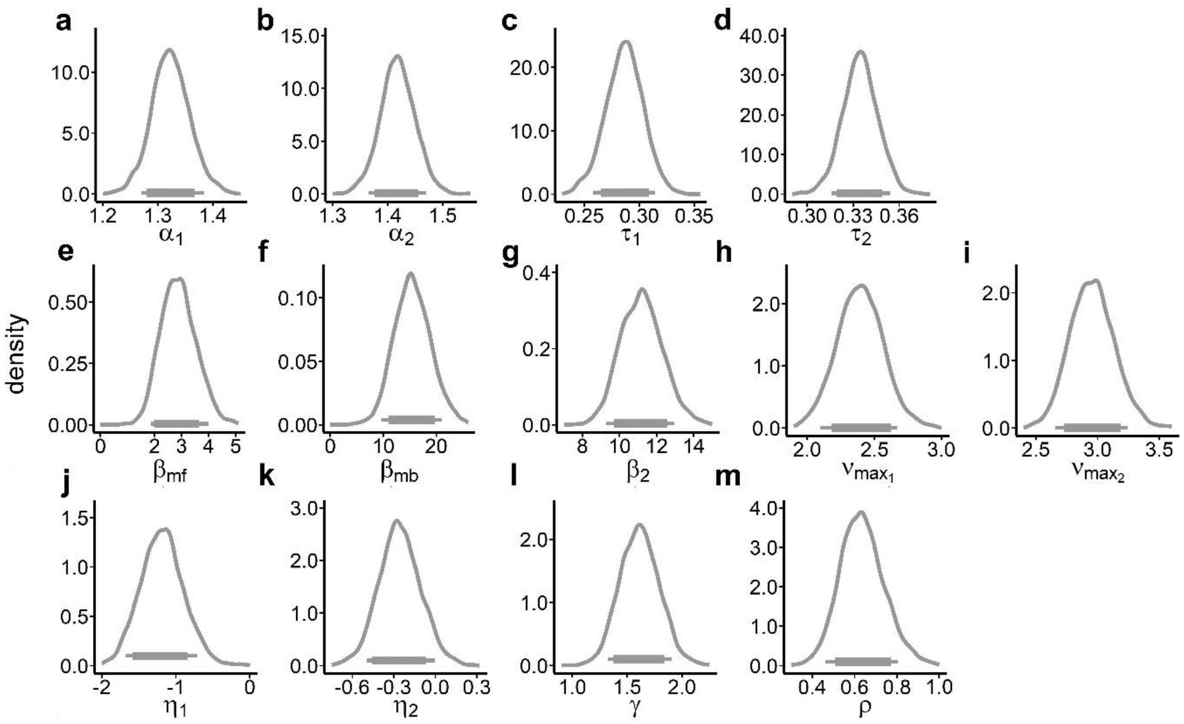


**Figure S3**. Group-level mean posterior distributions of the DDMs model parameter from the placebo condition for the seq. RL task data. Horizontal (thick) lines show 90 (80) % highest density intervals (HDIs). Depicted parameters are (a), (b) decision-thresholds $\alpha_1, \alpha_2$ (c), (d) non-decision times $\tau_1, \tau_2$; (e)-(g) weights for model-free Q-values $\beta_{mf}$, model-based Q-values $\beta_{mb}$ and S2 stage Q-values $\beta_2$; (h), (i) the asymptote of respective drift-rates $v_{max_1}$, $v_{max_2}$; (j), (k) learning-rates in S1 and S2 $\eta_1, \eta_2$, (l) decay rate of unchosen options $\gamma$ and (m) choice perseveration $\rho$.

*Sequential RL task, softmax model*

We also implemented an extension of an established RL model (Daw et al., 2011, Otto et al., 2015) that is based on softmax action selection instead of a DDM implementation (see Methods section). This model included separate parameters for S1 and S2 learning rates, a separate decay rate for unchosen options in both stages, model-free and model-based $\beta$ weights for S1 and a $\beta$ weight for S2 $Q$-value differences. Note that this model formulation includes the nested versions with one learning rate, no decay rate, no perseveration, no model-based and or no model-free effects as special cases, where the respective posteriors are centered at 0. Kruschke (2011) suggests to examine the posterior distributions of the full model in such cases, rather than performing a model comparison across all nested versions.

Similar to the mixed regression effects of reward and the reward*transition interaction above, we observed substantial contributions of both model-free and model-based values to S1 choice probabilities ($\beta_{MF}$, $\beta_{MB}$ in Figure S4 i, j). Participants also exhibited choice preservation in S1 and their choices in S2 were affected by S2 Q-value differences ($\rho$, $\beta_{S2}$ in Figure S4 d, k). Learning rates for updating of S1 and S2 model-free Q-values significantly differed (90% highest density interval (HDI) of $\eta_{S1}$ and $\eta_{S2}$ did not overlap, Figure S4 a, b) and unchosen choice option Q-values substantially decayed towards the mean ($\eta_{decay}$, Figure S4 c).

Tyrosine led to a significant decrease in the decay rate $\eta_{decay}$ in the softmax model (mean[90% HDI] = -.27 [-.48, -.08], directional Bayes Factor (dBF) = .01, Figure S4 g). According to dBF analysis, participants exhibited a substantial increase in updating of S1 model-free Q-values through heightened learning rates $\eta_{S1}$ (mean[90% HDI]=.54[-.07,1.09], dBF=16.3; Figure S4 e). All other posterior distributions of tyrosine-related parameter changes showed a substantial overlap with zero (80% HDIs, Figure S4).

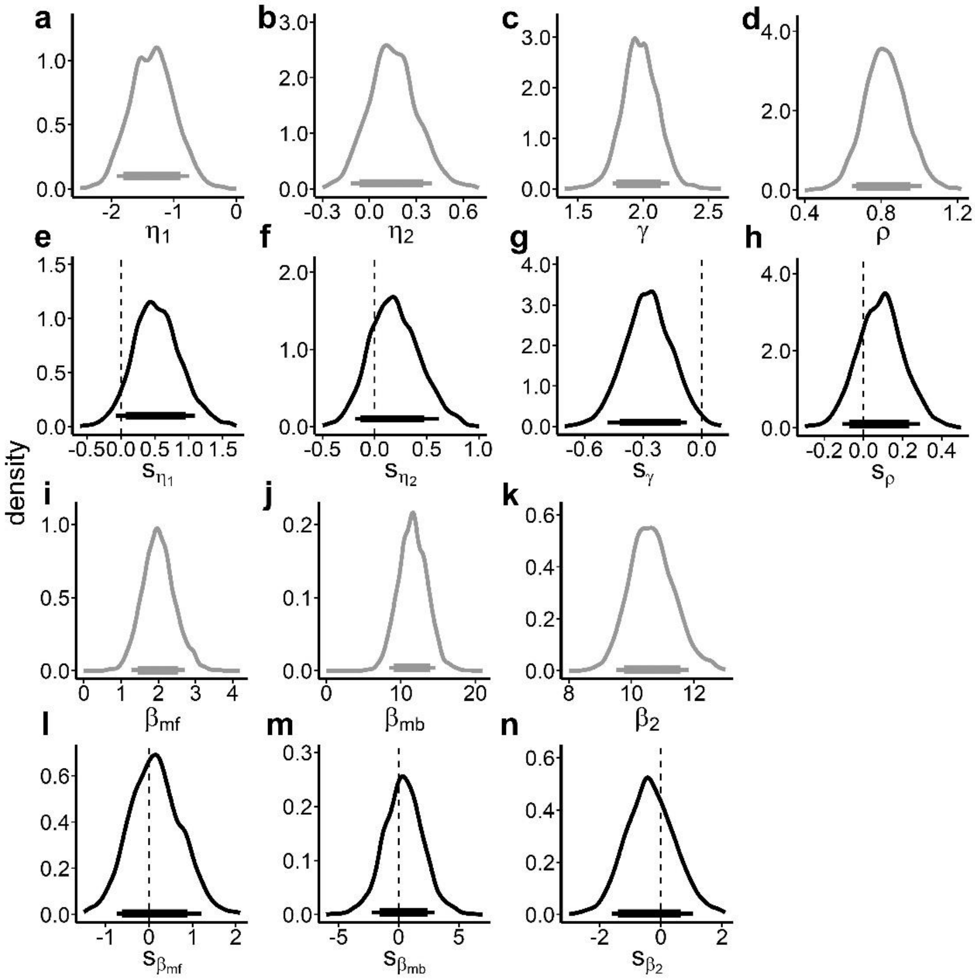


**Figure S4.** Posterior distributions of the group-level means of the softmax model of the seq. RL task data for the placebo condition (a-d, i-k; grey plots) and tyrosine related changes thereof (e-h, l-n; black plots). Horizontal (thick) lines show 90 (80) % HDIs. Depicted parameters are from left to right: (a) learning-rate in S1 $\eta_1$, (b) learning-rate in S2 $\eta_2$, (c) decay of unchosen options $\gamma$ , (d) choice perseveration $\rho$, (i) model-free $\beta_{mf}$ weight, (j) model-based $\beta_{mb}$ $weight,$ (k) S2 stage Q-value $\beta_2$ weight and their tyrosine related ‚shifts' ($s_x$ ; e-h, l-n; black plots).

*Temporal discounting task, softmax model*

We also modeled choice data in the temporal discounting task using a hyperbolic discounting model with standard softmax-action selection (see methods section). In this model, we found no evidence for a modulatory effect of tyrosine supplementation on discount rate log(k) (mean [90% HDI]: $k_s$=-.7 [-.28, .17], Figure S5). We also found no evidence for a modulatory effect of tyrosine on modeled choice stochasticity (mean [90% HDI]: $\beta_s$=.01 [-.04, .07], Figure S5).

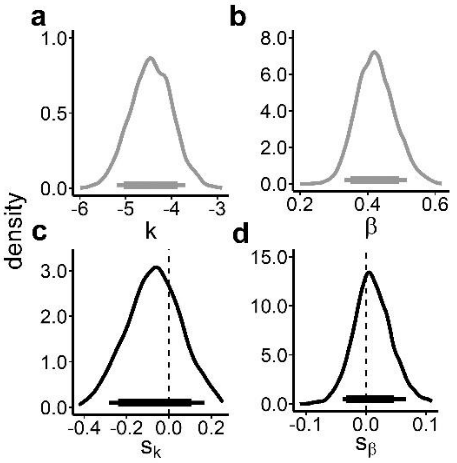


**Figure S5**. Posterior distributions of the group-level means of the softmax model of the temporal discounting task data for the placebo condition (a,b, grey plots) and tyrosine related changes thereof (c,d, black plots). Horizontal (thick) lines show 90 (80) % HDIs. Depicted parameters are: Discount rate $\log(k)$, softmax inverse temperature $\beta$, and tyrosine related shifts thereof $S_k$ , $S_\beta$ .

*Correspondence between task performance and physiology*

In an exploratory attempt, we tested for associations between task performance under placebo and physiological arousal measures (spontaneous eye blink rate, pupil dilation, pupil dilation variability, heart rate, heart rate variability; note that these values were averaged across placebo and tyrosine t0 baseline measurements), as well as tyrosine-related changes of these. We restricted these exploratory analyses to core aspects of seq. RL and temporal discounting, and related evidence accumulation processes to reduce multiple testing burden. For the seq. RL task, we focused on potential links between physiological measures and average payout per trial as an agnostic measure of general task performance, the impact of model-free and model-based Q-values on evidence accumulation speed ($\beta_{mf}$, $\beta_{mb}$), as well as participants' average (across task stages) decision thresholds $\alpha = mean(\alpha_1, \alpha_2)$ and non-decision times $\tau = mean(\tau_1, \tau_2)$. In addition, we tested for associations between tyrosine related changes in pupil dilation variability and heart rate (spontaneous eye blink rate, pupil dilation and heart rate variability were unaffected by tyrosine), and tyrosine associated shifts in decision thresholds $s_\alpha = mean(s_{\alpha_1}, s_{\alpha_2})$ and model-based control $s_{\beta_{mb}}$. Baseline pupil dilation was negatively associated with individual non-decision times during seq. RL ($\tau$; r=-.5, p=.007; Figure S6 a) and pupil dilation variability was positively correlated with the impact of model-free Q-values on trial-wise drift-rates ($\beta_{mf}$; r=.49, p=.009; Figure S6 b). All other tested associations were non-significant (all p>.14).

For testing potential associations between temporal discounting performance under placebo and physiological arousal measures, we focused on % LL choices as a model agnostic measure, discounting parameter log($k$), bias towards LL/SS choices $z$, decision thresholds $\alpha$, non-decision time $\tau$ and evidence accumulation speed $v$. In addition, we tested for associations between tyrosine related changes in pupil dilation variability and heart rate, and tyrosine associated shifts in discounting $s_{\log(k)}$ and decision thresholds $s_\alpha$. We found that higher pupil dilation at baseline was associated with more impatient (fewer LL) choices during temporal discounting (r=-.55, p=.002 ; Figure 7 a - main manuscript). In line, pupil dilation at baseline predicted steeper discounting (log($k$); r=.51, p=.005; Figure S6 c) and higher pupil dilation was associated with a significant bias towards impatient SS choices ($z$; r=-.63, p=3.0*10^-4; Figure 7 b – main manuscript). Higher pupil dilation variability at baseline was related to longer non-decision times ($\tau$; r=.46, p=.01). In addition, higher spontaneous eye blink rate at baseline was associated with lower decision thresholds during temporal discounting ($\alpha$; r=-.4, p=.03). With respect to tyrosine related changes in physiological arousal, greater pre-post heart rate changes following tyrosine compared with placebo were associated with tyrosine related shifts in temporal discounting ($s_{\log(k)}$; r=.61, p=5.76*10^-4; Figure 7 c – main manuscript). All other tested associations were non-significant (all p>.06).

However, when adjusting for False Discovery Rate (Benjamini & Hochberg, 1995) only the associations between pupil dilation and % LL choices, as well as bias towards SS choices, and between tyrosine related modulation of pre-post heart rate changes and tyrosine related shifts in temporal discounting remained significant (all other p-values >=.005; FDR adjusted p-value = .003).

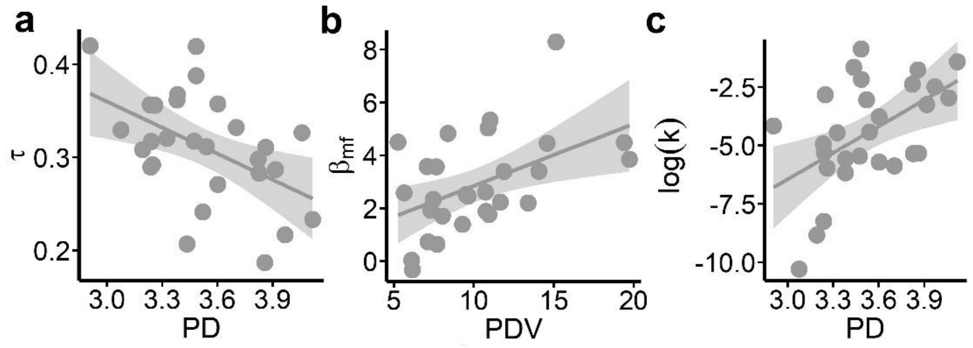


**Figure S6**. (a) Participants' pupil dilation at baseline (mean of t0 physiological measurements across tyrosine and placebo) was predictive of participants' average (S1 & S2) non-decision times ($\tau$) under placebo during seq (r=-.5, p=.007). RL. (b) Individual pupil dilation variability (PDV) at baseline was associated with the degree of drift-rate modulation by model-free Q-values ($\beta_{mf}$) during seq. RL under placebo (r=.49, p=.009). (c) Pupil dilation at baseline was also related to temporal discounting $\log(k)$ (r=.51, p=.005). Note, that the depicted associations fell short of significance after adjusting for False Discovery Rate (all p-values > FDR adjusted p-value = .003).
